# Allergen-induced dendritic cell migration is controlled through Substance P release by sensory neurons

**DOI:** 10.1101/2020.06.05.136929

**Authors:** Pamela A. Aderhold, Zaynah N. A. Dewan, Caroline Perner, Cameron H. Flayer, Xueping Zhu, Tiphaine Voisin, Ryan B. Camire, Isaac M. Chiu, Ohn A. Chow, Caroline L. Sokol

**Author notes:** These authors contributed equally to this work (Pamela A. Aderhold, Zaynah N. A. Dewan).

## Abstract

Dendritic cells (DCs) of the cDC2 lineage are necessary for the initiation of the allergic immune response and in the dermis are marked by their expression of CD301b. CD301b^+^ dermal DCs respond to allergens encountered in vivo, but not in vitro. This suggests that another cell in the dermis may sense allergens and relay that information to activate and induce the migration of CD301b^+^ DCs to the draining lymph node. Using a model of cutaneous allergen exposure, we show that allergens directly activate TRPV1^+^ sensory neurons leading to itch and pain behaviors. Allergen-activated sensory neurons release the neuropeptide Substance P, which stimulates proximally located CD301b^+^ DCs through MRGPRA1. Substance P induces CD301b^+^ DC migration to the draining lymph node where they initiate Th2 differentiation. Thus, sensory neurons act as primary sensors of allergens, linking exposure to activation of allergic-skewing DCs and the initiation of the allergic immune response.

## INTRODUCTION

Allergic diseases are characterized by inappropriate Type-2 immune responses targeted against non-infectious environmental antigens and venoms. Defective cutaneous barriers permit the entry of food and environmental allergens into the body where they can ultimately activate the immune system, but the precise mechanisms by which allergens are sensed by the innate immune system remain unclear (Strid et al., 2004). Dendritic cells (DCs) are innate immune cells that link pathogen sensing with adaptive immune activation. Immature DCs actively sample their environment for evidence of pathogens that are detected through their pattern recognition receptors (PRRs). Upon PRR ligation, DCs undergo a complex process of maturation that culminates in their migration to the draining lymph node (dLN), where they present antigen and provide costimulation to naïve T cells. Thus, DCs link innate immune sensing with adaptive immune activation.

Conventional DCs (cDC) can be divided into two main subsets based on transcription factor dependence and functional characteristics. cDC1s are dependent upon interferon regulatory factor 8 (IRF8) and are specialized in antigen cross-presentation, while cDC2s are dependent upon interferon regulatory factor 4 (IRF4) and are specialized in T helper cell differentiation. Th2-skewing cDC2s are dependent upon both IRF4 and Kruppel like factor 4 (KLF4), and in the skin this population is characterized by its surface expression of CD301b. CD301b^+^ DCs are required for Th2-differentiation after cutaneous exposure to allergens and helminth parasites and migrate to the dLN in response to this exposure (Gao et al., 2013; Kumamoto et al., 2013; Tussiwand et al., 2015). In the Type-1 immune response to bacteria and viruses, DC maturation is marked by the upregulation of CCR7. This CCR7 upregulation acts like a molecular switch, endowing the mature DC to sense and migrate towards gradients of CCL21 that are homeostatically produced by the lymphatic endothelium (Ohl et al., 2004). Although cutaneous allergen-exposure leads to CCR7-dependent CD301b^+^ DC migration from the skin to the dLN, it does not lead to CCR7 upregulation beyond the low levels expressed by immature DCs (Sokol et al., 2018). In the absence of this molecular switch, what is the primary signal that initiates CD301b^+^ DC migration out of the skin in response to allergens? Similarly, CD301b^+^ DCs are functionally activated by allergens in vivo and necessary to initiate Th2-differentiation in the dLN (Kumamoto et al., 2013; Sokol et al., 2018). However, CD301b^+^ DCs exposed to allergens in vitro are not capable of promoting Th2-differentiation from naïve CD4^+^ T cells in vitro or after in vivo transfer (Kumamoto et al., 2013). How do CD301b^+^ DCs sense allergens in vivo?

Type-2 immunogens are a varied group of simple protein allergens, secreted molecules from helminth parasites, and venoms (Palm et al., 2012). Many immunodominant Type-2 immunogens share in common intrinsic or induced enzymatic activity that is required for their immunogenicity, but how they activate the innate immune system is unclear (Lamhamedi-Cherradi et al., 2008; Palm et al., 2013; Porter et al., 2009; Sokol et al., 2008; Van Dyken and Locksley, 2018). It has been hypothesized that instead of direct detection, DCs may detect endogenous alarmins that are released in response to allergen exposure. Enzymatically active allergens such as cysteine proteases induce the release and activation of innate alarmins, such as IL-33 and thymic stromal lymphopoietin (TSLP) (Cayrol et al., 2018; Tang et al., 2010). Indeed, DCs express the IL-33 receptor ST2 and IL-33 stimulation of DCs can promote Th2 skewing (Rank et al., 2009). IL-33 can also stimulate innate lymphoid cells, which can indirectly enhance DC migration to dLNs (Halim et al., 2014). However, data from alarmin-deficient models show only a partial role for alarmins in allergen-induced DC migration and activation (Besnard et al., 2011; Halim et al., 2014). This led us to hypothesize the existence of a primary cellular sensor of allergens that not only senses the presence of allergens, but also relays critical cues to initiate the migration of allergic, or Th2-skewing, CD301b^+^ DCs to the dLN.

Mast cell degranulation, a process that classically occurs when allergens cross link allergen-specific IgE, specifically promotes the migration of Th2-skewing DCs to the dLN (Mazzoni et al., 2006). DCs have been shown to transfer antigens to mast cells leading to degranulation, which then can feed back and enhance DC migration (Choi et al., 2018). These pathways, which largely depend on the presence of antigen-specific IgE, may play important roles in amplifying or boosting allergic memory responses. However, in the naïve animal we hypothesized that a non-immune cell may be the initial sensor of allergens.

One potential allergen sensor is the sensory nervous system, which is highly concentrated in barrier epithelia, broadly responsive to many different stimuli, and directly activated in vitro by diverse enzymatically active allergens such as bee venom, house dust mites, and papain (Chen and Lariviere, 2010; Reddy and Lerner, 2010; Serhan et al., 2019; Talbot et al., 2016; Trier et al., 2019; Veiga-Fernandes and Mucida, 2016). Sensory neurons have been shown to be closely associated with Langerhans cells in the human epidermis and activation of corneal sensory neurons can lead to DC activation in the contralateral eye (Guzman et al., 2018; Hosoi et al., 1993). Based on this, we hypothesized a two-step model for allergen activation of DCs wherein allergens activate sensory neurons leading to their local release of neuropeptides (Step 1), which then act upon local DCs to promote their migration (Step 2) to the dLN where they can activate naïve T cells to promote Th2 differentiation. Using the model cysteine protease allergen papain, we found that papain induces an immediate and transient sensory response in naïve mice. Papain directly activates a subset of sensory neurons that are enriched in TRPV1^+^ fibers to release Substance P. Substance P then acts on proximally located Th2-skewing dermal CD301b^+^ DCs to promote their migration to the dLN. Inhibition of sensory neuronal activation through chemical inhibitor or depletion of TRPV1^+^ neurons leads to a defect in CD301b^+^ DC migration and as a direct consequence, Th2-differentiation. Thus, sensory neurons play an essential role in allergen recognition, DC activation, and initiation of the allergic immune response.

## RESULTS

### Allergens induce immediate and transient itch responses in naïve mice

To investigate the role of sensory neurons in allergen recognition in vivo, we utilized the model allergen papain that induces robust CD301b^+^ DC migration, Th2 differentiation and IgE production with defined kinetics after one cutaneous exposure to the enzymatically active allergen (Kumamoto et al., 2013; Sokol et al., 2008). Intradermal papain (i.d.) injection of mice led to an immediate and transient mixed itch (scratching bouts) and pain (wiping bouts) response that was dependent on its protease activity (Fig. 1A-D). Papain induced an itch response that was faster in onset, but overall comparable to the known itch trigger histamine (Fig. 1A-D). Likewise, the papain-induced pain response was comparable to the known pain trigger capsaicin (Fig. 1A-D). The sensory response was not specific to papain as Alternaria extract was similarly capable of inducing an itch response (Fig. 1A & B). Mast cells have been shown to play a role in amplifying sensory neuron activation and tissue inflammation by house dust mite (Serhan et al., 2019). To investigate whether mast cells were similarly involved in the immediate sensory response to papain, we injected papain i.d. in wild type or mast cell deficient *Kit^W-sh/W-sh^* mice. The sensory response to papain was unaffected by mast cell deficiency indicating that mast cells are not required for this sensory response (Fig. 1E). However, we found that the sensory response was inhibited by papain co-injection with the lidocaine derivative QX314 (Fig. 1F). QX314 blocks sodium channel activation of neurons after entering through large-pore cation channels including TRPV1 and TRPA1 (Binshtok et al., 2007; Roberson et al., 2013). Thus, the inhibition of the papain-induced itch and pain response suggested a role for TRPV1^+^ neurons in direct allergen sensing.

**Figure 1.**
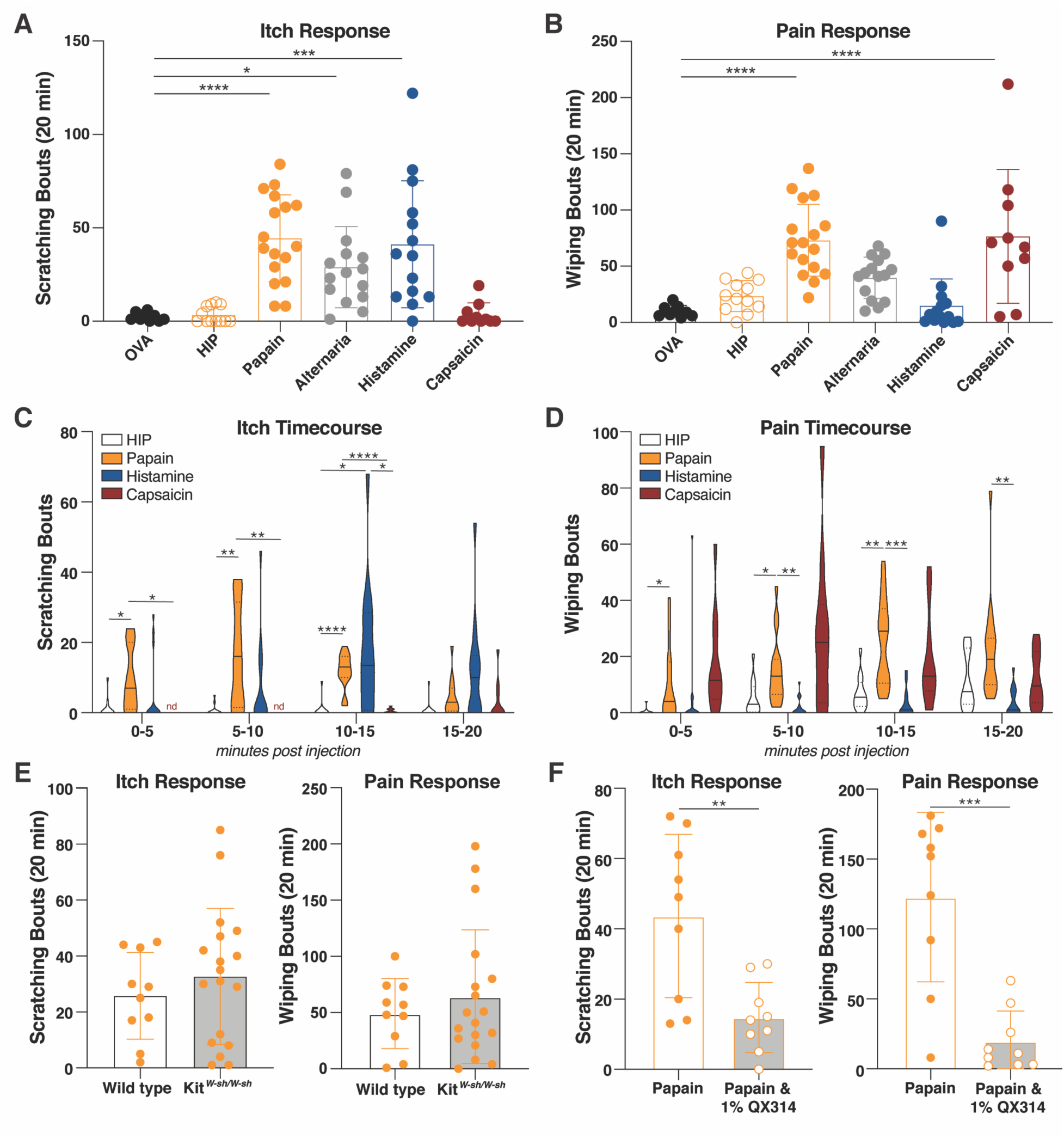
The protease allergen papain induces a mast cell independent sensory response. (**A** and **B**) Wild type (C57Bl/6) mice were intradermally (i.d.) injected with ovalbumin (OVA), heat inactivated papain (HIP), papain, Alternaria extract, histamine, or capsaicin. The total number of ipsilateral (A) cheek scratching events or (B) cheek wiping events were counted in the 20 minutes after i.d. cheek injection. (**C** and **D**) Timecourse of (C) cheek scratching or (D) cheek wiping events in 5 minute increments from mice shown in (A and B). (**E**) Wild type (white histogram) or mast cell-deficient *Kit^w- sh/w-sh^* (gray shaded histogram) mice were i.d. injected with papain in the cheek and total number of cheek scratching or wiping events were counted in the 20 minutes after injection. (**F**) Wild type mice were i.d. cheek injected with papain or papain & 1% QX314. The total number of cheek scratching and wiping events in the 20 minutes after injection is shown. Each symbol represents an individual mouse (A, B, E, F), bar indicates mean, and error bars indicate SEM. Violin plots (C and D) show grouped timecourse data from mice (A and B); line indicates median, dotted lines indicate quartiles, and nd indicates not detected. Statistical tests: Ordinary one-way ANOVA with multiple comparisons (A and B), two-way ANOVA with multiple comparisons (C and D), and unpaired t test (E and F). * p<0.05, ** p<0.01, *** p<0.001, **** p<0.0001. Data are representative of at least three independent experiments combined with each experiment including 3-5 mice per group.

### Allergens directly activate a population of sensory neurons enriched in TRPV1^+^ fibers

The cell bodies of cutaneous sensory neurons are contained in dorsal root ganglia (DRG) structures adjacent to the spinal cord. To test whether papain could directly activate sensory neurons, we performed calcium (Ca^2+^) flux analysis of cultured DRG neurons. Papain induced robust Ca^2+^ influx in DRG neurons within seconds of administration (Fig. 2A & B). Using potassium chloride (KCl) responsiveness as a marker of live neurons, we found that approximately 16% neurons were responsive to papain (Fig. 2C), a similar value to that reported for house dust mite responsive neurons (Serhan et al., 2019). TRPV1 expression defines a subset of nociceptive neurons that not only sense noxious heat, but also promote pruriceptive responses. These TRPV1^+^ neurons are directly activated by house dust mite cysteine protease allergens and have been associated with allergic inflammation (Roberson et al., 2013; Serhan et al., 2019; Talbot et al., 2015; Trankner et al., 2014). Papain-responsive neurons were largely TRVP1^+^ (capsaicin-responsive) neurons, with an average of 68% of papain-responsive neurons also responding to capsaicin (Fig. 2D & E). A minority of papain-responsive neurons were also activated by the itch inducing ligands histamine and/or chloroquine, suggesting that papain-responsive neurons are a heterogenous population (Fig. 2D & E). Given that papain-responsive neurons strongly overlapped with TRPV1^+^ (capsaicin-responsive) neurons, we next determined whether the papain-induced sensory response would be affected by depletion of noxious heat-sensing TRPV1^+^ neurons. We used *Trpv1^DTR^* mice, which express the human diphtheria toxin receptor (DTR) under control of the *Trpv1* promoter (Pogorzala et al., 2013). Diphtheria toxin (DT) injection into *Trpv1^DTR^* mice has been shown to specifically deplete TRPV1^+^ neurons in the DRG and vagal ganglia (Baral et al., 2018; Trankner et al., 2014). Consistent with depletion of TRPV1^+^ neurons, DT treated *Trpv1^DTR^* mice lost their tail flick response to noxious heat and *Trpv1* gene expression in the dorsal root ganglia (DRG) (Fig. 2 F **and** G). Deletion of *Trpv1*^+^ neurons led to a loss of the itch response and a partial block in the pain response to papain (Fig. 2 H **and** I), indicating that the cysteine protease allergen papain activates TRPV1^+^ neurons in vitro and in vivo to induce an immediate itch response.

**Figure 2.**
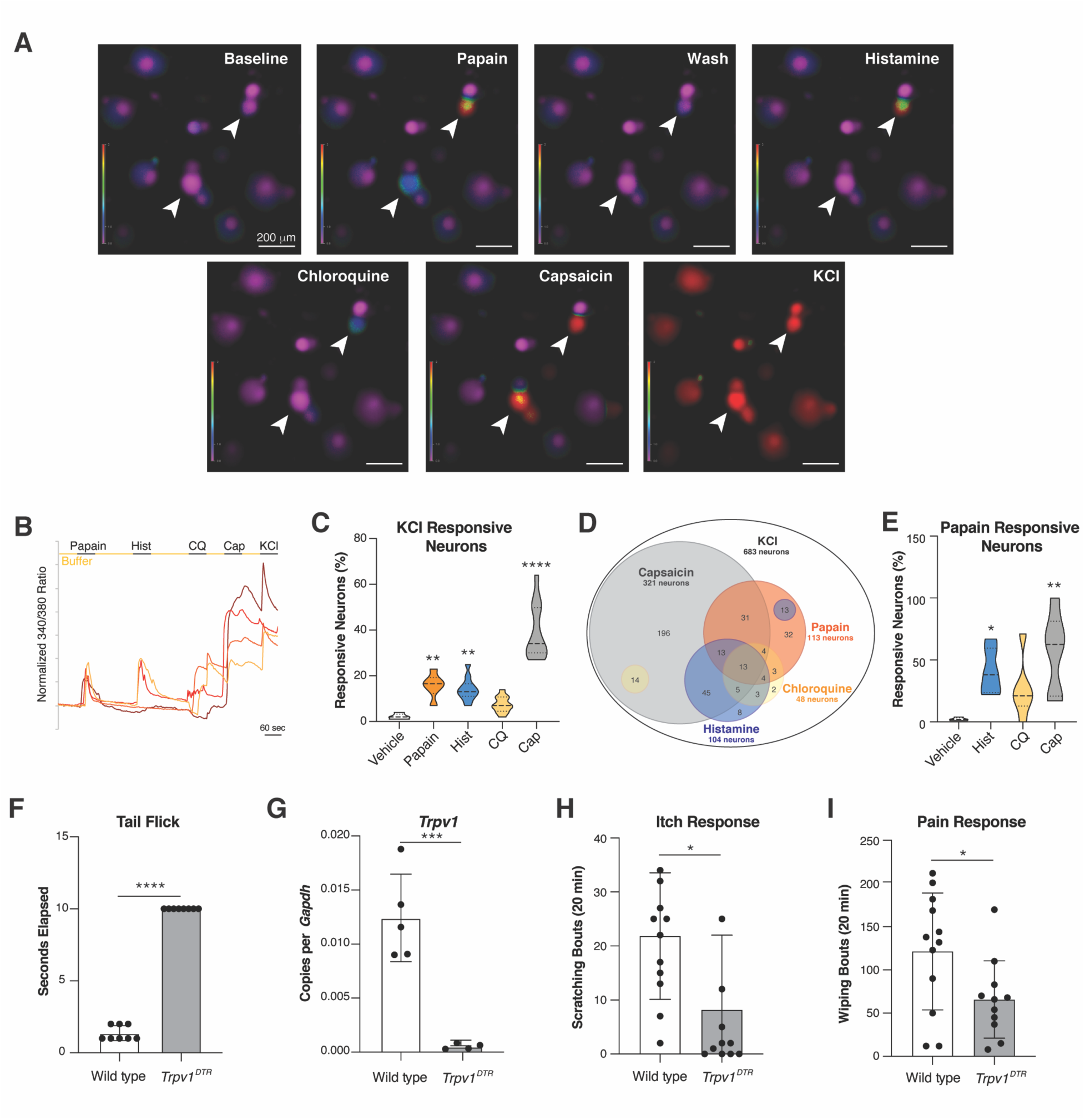
TRPV1^+^ neurons are directly activated by papain and are required for the papain-induced sensory response. (**A**) Representative Fura-2-AM ratiometric fields of cultured dorsal root ganglia (DRG) neurons sequentially treated with vehicle (Baseline), 100 µg/ml papain, vehicle (Wash), 100 µM Histamine, vehicle, 1 mM chloroquine, vehicle, 1 µM capsaicin, vehicle, and 80 mM KCI. (**B**) Calcium traces of representative Fura-2-AM loaded individual DRG neurons treated as in (A). Each neuron is colored separately to allow identification. (**C**) The percent of total (KCl responsive) DRG neurons that respond to vehicle, papain, histamine, chloroquine, or capsaicin. Neurons were calculated from 12 fields of view and a total of 1389 neurons, with 6 fields of view and 705 neurons for papain responsiveness and 684 neurons for vehicle responsiveness. (**D**) Venn diagram showing overlap in responsiveness of individual DRG neurons sequentially stimulated with papain, histamine, chloroquine, and capsaicin out of total (KCl responsive) neurons. Numbers of neurons in each group shown out of a total of 683 KCl responsive, but vehicle non-responsive neurons calculated from 6 fields of view and a total of 705 neurons. (**E**) The percent of papain responsive DRG neurons that also responded to histamine, chloroquine, or capsaicin were calculated from 6 views with a total of 705 neurons. This is compared to the percent of vehicle responsive neurons counted from 6 fields of view with a total of 684 neurons investigated. (**F**) Wild type or *Trpv1^DTR^* mice were intraperitoneally injected with diphtheria toxin (DT) for 21 days and rested for 7 days before undergoing a tail flick assay. Tails of restrained mice were applied to 52°C water and the time before the mouse removed its tail (“tail flick”) was recorded. A maximum water exposure of 10 seconds was allowed before removal. (**G**) DRGs were removed from wild type or *Trpv1^DTR^* mice treated with DT as in (F). Copies of *Trpv1* normalized to copies of *Gapdh* are shown. (**H** and **I**) Wild type or *Trpv1^DTR^* mice treated as in (F) were i.d. cheek injected with papain and the total number of cheek scratching (H) and cheek wiping (I) events were counted in the 20 minutes after injection. Each symbol represents an individual mouse (F, G, H, I), bar indicates mean, and error bars indicate SEM. Violin plots (C and E) show neuron responsiveness to different stimuli as measured by Ca^2+^ flux, line indicates median, dotted lines indicate quartiles. Statistical tests: Ordinary on-way ANOVA with multiple comparisons (C and E), unpaired t test (F, G, H, I). * p<0.05, ** p<0.01, *** p<0.001, **** p<0.0001. Data are representative of at least three independent experiments combined with each experiment including 3-5 mice per group.

### Dendritic cells do not directly respond to the cysteine protease allergen papain

DCs directly detect bacterial and viral antigens through their expression of PRRs. Pathogen associated molecular patterns (PAMPs) directly bind to PRRs and lead to DC maturation, marked by the upregulation of antigen presentation, costimulatory molecules, and CCR7. Thus, maturation promotes the migration of DCs with unique T cell activating capability to the dLN. We performed flow cytometry of DCs from the dLN 24 hours after in vivo immunization with lipopolysaccharide (LPS) and after in vitro exposure to LPS. Consistent with the direct activation of DCs by PAMPs, CD301b^+^ DCs exposed to LPS in vivo or in vitro upregulate the activation marker PDL2 (Fig. 3A **and** B). Whereas in vivo papain immunization was able to induce PDL2 upregulation, in vitro papain stimulation had no effect on PDL2 expression on CD301b^+^ DCs (Fig. 3A **and** B). Similarly, in vivo immunization with papain and LPS led to an increase in the chemotactic activity of CD301b^+^ DCs (Fig. 3C), while in vitro exposure with papain had no effect (Fig. 3D). These observations suggest that DCs do not independently detect allergens.

**Figure 3.**
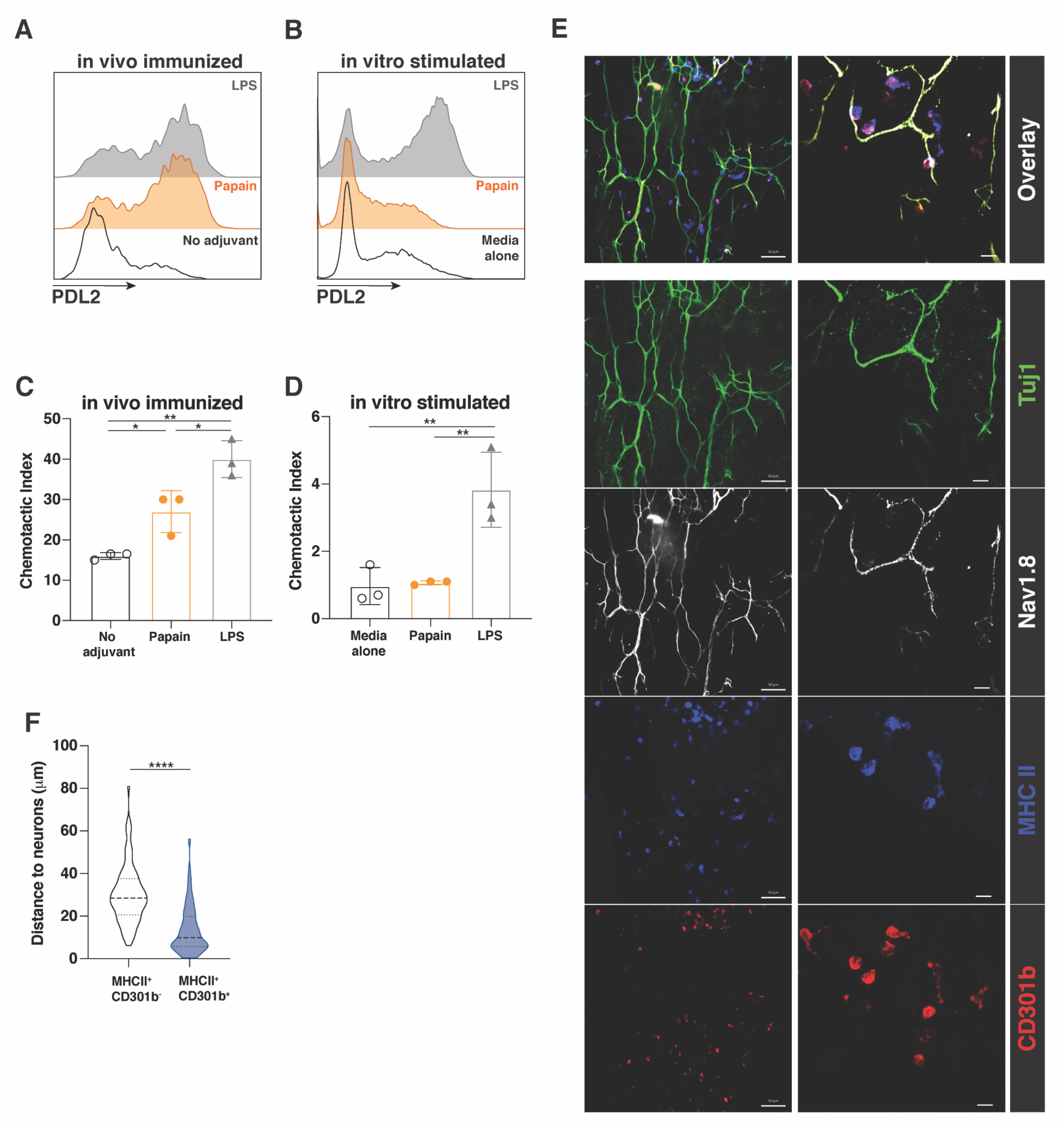
CD301b^+^ dendritic cells do not respond directly to the protease allergen papain and are found in close proximity to sensory neurons in the dermis. (**A**) Mice were immunized i.d. with ovalbumin (OVA) in PBS (No adjuvant), OVA and papain (Papain), or OVA and lipopolysaccharide (LPS). Flow cytometric expression of the activation marker PDL2 on live CD11c^+^CD301b^+^ dendritic cells (DCs) was determined 24 hours after immunization. (**B**) Bone marrow derived DCs (BMDCs) were stimulated overnight with media alone, papain, or LPS, with the expression of PDL2 on CD11c^+^CD301b^+^ BMDCs being measured by flow cytometry. (**C** and **D**) Transwell migration to CCL21 of (C) CD11c^+^CD301b^+^ cells sorted from the draining lymph node (dLN) of mice immunized as in (A) or of (D) BMDCs stimulated as in (B). Chemotactic index in C and D shows relative migration of cells to CCL21 in comparison to media only control. (**E**) Confocal immunofluorescence microscopy of unimmunized dermal sheets from *Nav1.8^mCherry^* (shown in white) mice stained for the pan-neuronal marker Tuj1 (green), MHCII (blue) and CD301b (red). Scale bar in top 20x panels shows 50 μm, scale bar in bottom 63x panels shows 10 μm. (**F**) Closest distance (μm) between MHCII^+^CD301b^-^ (clear) and MHCII^+^CD301b^+^ (blue) cells and mCherry^+^ neurons from naïve *Nav1.8^mCherry^* mice. Symbols represent individual replicates (C and D). Bars indicate mean and error bars indicate SEM. Violin plot (F) shows distance (μM) between cells and neurons, 10 fields of view were counted and 81 MHCII^+^CD301b^-^ cells and 122 MHCII^+^CD301b^+^ cells were counted, line on violin plot indicates median, dotted lines indicate quartiles. Statistical tests: Ordinary one-way ANOVA with multiple comparisons (C and D), unpaired t test (F). * p<0.05, ** p<0.01, *** p<0.001, **** p<0.0001. Data are representative of at least three experiments (A-F), combined in F with each experiment including 2-5 mice per group.

### CD301b^+^ dendritic cells colocalize with sensory neurons

Given our observations that sensory neurons, but not DCs, are directly activated by allergens, we hypothesized that allergens are detected by sensory neurons that then relay this signal to local DCs. To determine whether sensory neurons could be interacting with CD301b^+^ DCs in vivo, we imaged the naïve skin of *Nav1.8*^Cre/+^*tdTomato^loxSTOPlox^* reporter mice where *Nav1.8*-lineage nociceptive sensory neurons are marked by tdTomato expression (Stirling et al., 2005). CD301b^+^ DCs were visualized in close proximity to the Nav1.8^+^ nerve fibers in the naïve dermis (Fig. 3E). Furthermore, CD301b^+^ DCs were significantly closer to sensory neurons than were their CD301b- counterparts, which in the dermis include the Th1-skewing cDC1 and the Th17-skewing subset of cDC2 (Fig. 3F). Given these observations as well as published data showing evidence for sensory neuron interactions with CD301b^+^ DCs in the context of Candida infection (Kashem et al., 2015), we hypothesized that the relative proximity of Th2-skewing DCs to sensory neurons could reflect a functional requirement for neuronal stimulation specific to allergen-responsive CD301b^+^ DCs in the dermis.

### TRPV1^+^ sensory neuron activation is required for allergen-induced CD301b^+^ DC migration

We hypothesized that sensory neurons, activated by allergens, may relay a signal to closely associated CD301b^+^ DCs to activate their allergen-induced migration to the dLN. To assess allergen-induced migration, we used Kaede mice that express a photoconvertible green to red fluorescent protein, which allows the tracking of cells originating from the site of photoconversion and allergen exposure (Kaede^red^) (Tomura et al., 2008). As previously described, papain immunization led to a specific migration of CD301b^+^ DCs from the photoconverted skin to the dLN (Fig. 4A) (Sokol et al., 2018). QX314 co-injection blocked this papain-induced migration of skin emigrant (Kaede^red^) CD301b^+^ DCs to the dLN (Fig. 4A), suggesting a role for sensory neuron activation in CD301b^+^ DC migration in vivo. To evaluate for possible DC-specific effects of QX314 we performed in vitro chemotaxis assays of bone marrow derived DCs (BMDCs). QX314 had no effect on BMDC migration in vitro, indicating a specific role for neurons in allergen-induced DC migration (Fig. 4B).

**Figure 4.**
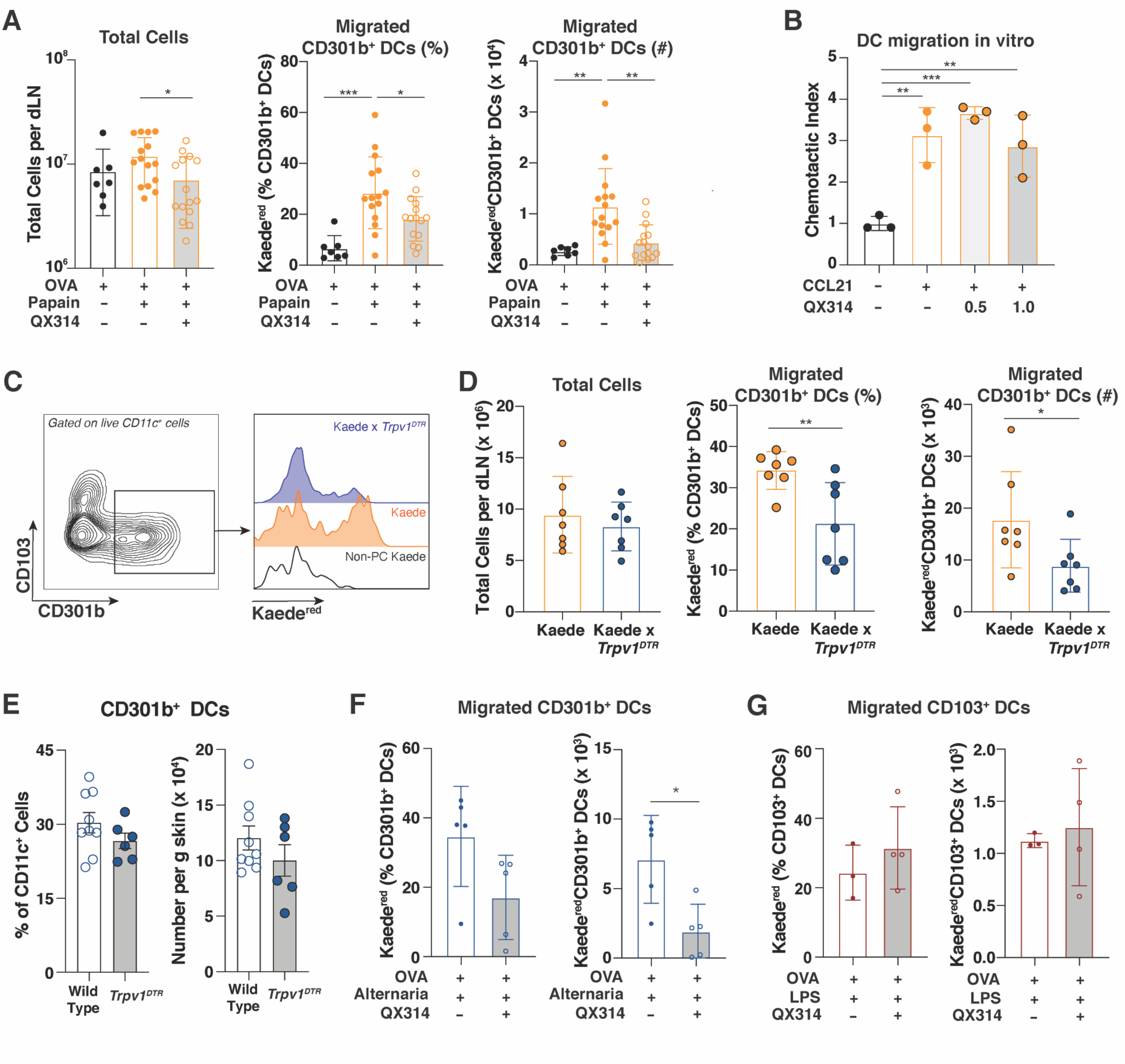
Allergen-induced migration of cutaneous Th2-skewing CD301b^+^ dendritic cells into the draining lymph node requires TRPV1^+^ sensory neurons. (A) The base of tail skin of *Kaede* transgenic mice was photoconverted and then immunized intradermally with the indicated combinations of OVA, papain and 1% QX314. 24 hours after immunization the dLN was harvested and examined for total cell number and the number and the percentage of CD11c^+^CD301b^+^ cells (CD301b^+^ DCs) that migrated from the skin (Kaede^red^). (**B**) BMDCs were stimulated overnight with LPS and placed on 5 μm Transwell membrane. Migration to 100 ng/ml CCL21 in the presence or absence of QX314 over a 2 hour incubation is shown. (**C**) *Kaede* or *Kaede* x *Trpv1^DTR^* mice were treated with DT for 21 days followed by 7 days of rest. Mice were then photoconverted, i.d. immunized with papain, and the dLN was harvested for flow cytometry. Flow cytometry of CD103 and CD301b expression on live CD11c^+^ cells and histogram of Kaede^red^ expression on CD301b^+^ DCs from the indicated gate are shown (black histogram indicates non-photoconverted control, orange histogram indicates *Kaede* mouse, blue histogram indicates *Kaede* x *Trpv1^DTR^* mouse). (**D**) *Kaede* and *Kaede* x *Trpv1^DTR^* mice were treated as in (C) and total cells per dLN, the percentage of skin emigrant CD301b^+^ DCs (Kaede^red^), and the total number of Kaede^red^CD301b^+^ DCs was quantified by flow cytometry. (**E**) Percent of MHCII^+^CD301b^+^ DCs out of total CD45^+^CD11c^+^ cells in the dermis or number of CD11c^+^MHCII^+^CD301b^+^ cells per gram (g) skin in wild type or *Trpv1^DTR^* mice treated with DT for 21 days followed by 7 days of rest. (**F**) *Kaede* mice were photoconverted as in (A) and immunized with OVA and Alternaria extract in the presence or absence of 1% QX314. Flow cytometry of Kaede^red^ expression on CD11c^+^CD301b^+^ (CD301b^+^ DCs) was performed, with the percent of Kaede^red^ cells out of total CD301b^+^ DCs or number is shown. (**G**) Kaede mice were photoconverted as in (A) and immunized with OVA and LPS in the presence or absence of 1% QX314. Flow cytometry of Kaede^red^ CD103^+^ DCs as a percent of total CD103^+^ DCs or number is shown. Symbols represent individual mice (A, D-G) or replicates (B). Bars indicate mean and error bars indicate SEM. Statistical tests: Ordinary one-way ANOVA with multiple comparisons (A and B), unpaired t test (D-G). * p<0.05, ** p<0.01, *** p<0.001, **** p<0.0001. Data are representative of at least three (A-D) or two (E-G) independent experiments, combined in A, D, and E with each experiment including 2-5 (A and D) or 5-9 (E) mice per group.

Given our data that papain activated a population of neurons enriched in TRPV1^+^ sensory neurons (Fig. 2D **and** E), and that mice depleted of TRPV1^+^ neurons exhibited defective sensory responses to papain (Fig. 2H **and** I), we hypothesized that TRPV1^+^ neurons would be required for papain-induced CD301b^+^ DC migration. To specifically assess DC migration from the skin, we generated Kaede x *Trpv1^DTR^* mice and performed DT-mediated depletion of Trpv1^+^ neurons. Papain-immunized Kaede x *Trpv1^DTR^* mice showed a loss of Kaede^red^ (skin emigrant) CD301b^+^ DCs compared with Kaede mice (Fig. 4 C). Although QX314 injected mice showed a mild decrease in cellular hypertrophy of the dLN (Fig. 4 A), TRPV1-depleted mice showed no decrease in dLN cellularity (Fig. 4 D). However, TRPV1-depleted Kaede x *Trpv1^DTR^* mice exhibited significantly decreased migration of CD301b^+^ DCs from the skin to the dLN (Fig. 4 D). The absence of TRPV1^+^ neurons in DT treated *Trpv1^DTR^* mice had no effect on the overall frequency or number of CD301b^+^ DCs in the naïve dermis (Fig. 4E). This observation suggests that disrupted CD301b^+^ DC migration to the dLN following papain immunization in *Trpv1^DTR^* mice was due to a specific defect in migration, not homeostatic reduction in cell number. Like papain, Alternaria extract led to CD301b^+^ DC migration from the skin to the dLN that was blocked by QX314 indicating that sensory neuron activation may be a global requirement for allergen-induced CD301b^+^ DC migration (Fig. 4F). Finally, this requirement for sensory neuron activation was specific to allergen-induced CD301b^+^ DC migration. There was no effect of QX314 on the migration of CD103^+^ DCs (including cDC1 and Th17-skewing cDC2s in the dermis) after immunization with OVA and LPS (Fig. 4G).

### Substance P release from sensory neurons induces CD301b^+^ DC migration

Our data suggested that TRPV1^+^ sensory neurons directly sense cysteine protease allergens and relay signals to promote the egress of local Th2-skewing CD301b^+^ DCs to the dLN. Activated peptidergic TRPV1^+^ neurons can release Substance P in response to allergen exposure and CGRP in response to bacterial and fungal exposures (Baral et al., 2018; Chiu et al., 2013; Kashem et al., 2015; Pinho-Ribeiro et al., 2018; Serhan et al., 2019). CGRP release was shown to promote IL-23 secretion by CD301b^+^ DCs after infection with the fungus *Candida* (Cohen et al., 2019; Kashem et al., 2015). Papain immunization promoted Substance P and inhibited CGRP release from skin explants (Fig. 5A). This pattern of neuropeptide release was also seen after direct stimulation of DRG cultures with both papain and house dust mite (HDM) extract (Fig. 5B **and S1**), indicating that cysteine protease allergens directly stimulate neuropeptide release from sensory neurons. In contrast to another report, we found that Alternaria extract, which has serine protease activity, was also capable of inducing DRG release of Substance P and inhibition of CGRP (**Fig. S1**) (Cayrol et al., 2018; Serhan et al., 2019). These data indicate that the capacity for sensory neuron activation and Substance P release may be shared amongst disparate allergens.

**Figure 5.**
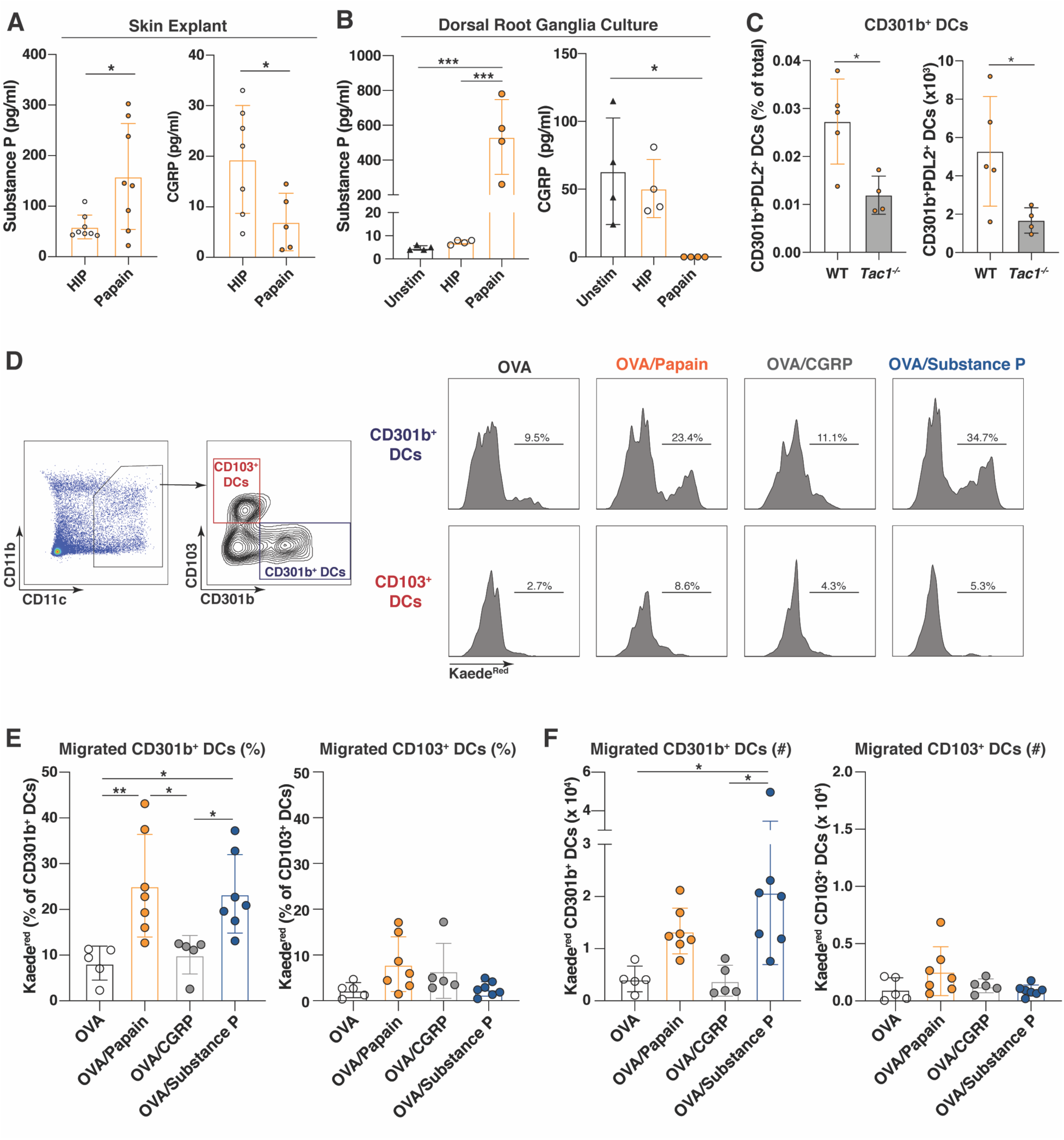
Allergen-stimulated sensory neurons release Substance P, which induces the specific migration of Th2-skewing CD301b^+^ dendritic cells into the draining lymph node. (**A**) Wild type mice were i.d. injected with heat inactivated papain (HIP) or papain, skin explants of the injected site were harvested and incubated in serum free media, and supernatant was tested by ELISA for Substance P and CGRP. (A) Dorsal root ganglia were harvested from wild type mice and were left unstimulated with PBS, or stimulated with either HIP or papain. Supernatants were measured by ELISA for Substance P and CGRP release. (**C**) Wild type (WT) or *Tac1^-/-^* mice were i.d. immunized with OVA/Papain and the dLN was harvested 24 hours later to evaluate the percent of activated (PDL2^+^) CD301b^+^ DCs out of total lymph node cells as well as the total number of activated PDL2^+^CD301b^+^ DCs per draining lymph node. (**D**) The skin of *Kaede* mice was photoconverted and i.d. immunized with OVA, OVA/Papain, OVA/CGRP or OVA/Substance P. The dLN was harvested 24 hours after immunization and total percent of skin emigrant Kaede^red^ cells was determined by flow cytometry. Flow cytometry gating scheme of photoconverted and immunized *Kaede* mice, showing the gates for CD11c^+^ DCs, followed by CD103^+^ and CD301b^+^ DCs. Percentage of Kaede^red^CD301b^+^ DCs and Kaede^red^CD103^+^ DCs in response to each immunization. (**E** and **F**) The (E) percentage and (F) total number of CD301b^+^ DCs and CD103^+^ DCs that originated from the skin of *Kaede* mice photoconverted and immunized as in (D). Symbols represent individual mice (A, C, E, F) or replicates (B). Bars indicate mean and error bars indicate SEM. Statistical tests: unpaired t test (A, C), ordinary one-way ANOVA with multiple comparisons (B, E, F). * p<0.05, ** p<0.01, *** p<0.001. Data are representative of at least three independent experiments (B, C, D), combined in (A, E, F), with each experiment including 2-4 mice per group.

We next examined the role of Substance P in promoting CD301b^+^ DC migration. Substance P deficient *Tac1^-/-^* mice showed a decrease in the percent and total number of activated (PDL2^+^) CD301b^+^ DCs in the dLN 24 hours after papain immunization (Fig. 5C). To specifically assess the role of Substance P and CGRP in CD301b^+^ DC migration from the skin, we injected photoconverted skin of Kaede mice with OVA and CGRP or Substance P. Although CGRP had no significant effect on CD301b^+^ or CD103^+^ DC migration, Substance P injection robustly induced the migration of Kaede^red^ CD301b^+^ DCs to the dLN (Fig. 5D-F). This effect of Substance P was specific to the migration of Th2-skewing CD301b^+^ DCs; Th1 and Th17-skewing CD103^+^ dermal DCs were unaffected by Substance P injection (Fig 5D-F). Following allergen immunization, Substance P is necessary and sufficient to induce the migration of Th2-skewing CD301b^+^ DCs to the dLN, suggesting that diverse allergens may induce CD301b^+^ DC migration in a consistent manner.

### Allergen-induced Substance P release and CD301b^+^ DC migration is mast cell independent

Mast cells and sensory neurons interact and can promote each other’s function in a positive-feedback manner. Substance P, released from sensory neurons, can activate mast cells through the receptor MRGPRB2 leading to mast cell degranulation, inflammation, and pain (Green et al., 2019; Serhan et al., 2019). At the same time, Substance P can directly activate sensory neurons, potentially boosting any response induced by direct neuronal activation (Azimi et al., 2017). Conversely, mast cell degranulation products such as trypsin can activate local sensory neurons leading to non-histaminergic itch (Meixiong et al., 2019a). To investigate the role of neuronal versus mast cell function in allergen-induced Substance P release, we first examined whether inhibiting neuronal activation altered Substance P release. Co-injection of papain with QX314 led to an incomplete, but significant decrease in Substance P release from flank explants indicating a major role for neuronal depolarization in allergen-induced Substance P release (Fig. 6A). To assess whether local mast cells played a role in a positive feedback loop with neurons, we examined Substance P release from wild type versus mast cell deficient *Kit^W-sh/W-sh^* mice. In response to papain injection, mast cell deficient mice were capable of robust Substance P release from skin explants with no significant difference noted from wild type controls (Fig. 6 B). These data indicate that neuronal activation, but not mast cell activation, is required for Substance P release. Mast cells have been shown to promote DC migration, although not all DC subsets appear to be affected equally (Dawicki et al., 2010; Shelburne et al., 2009). Since Substance P was shown to activate mast cells we next asked whether mast cells were involved in the migration of CD301b^+^ DCs to the dLN in response to papain. Wild type and mast cell deficient *Kit^W-sh/W-sh^* mice were immunized and their dLNs were examined 24 hours later to assess for CD301b^+^ DC migration by examining the quantity of activated (PDL2^+^) CD301b^+^ DCs in the dLN. We found no difference in the frequency or number of CD301b^+^PDL2^+^ DCs in dLNs after papain immunization (Fig. 6C). Thus, CD301b^+^ DC migration in response to allergen in naïve mice is independent of mast cells, but dependent on TRPV1^+^ sensory neurons.

**Figure 6.**
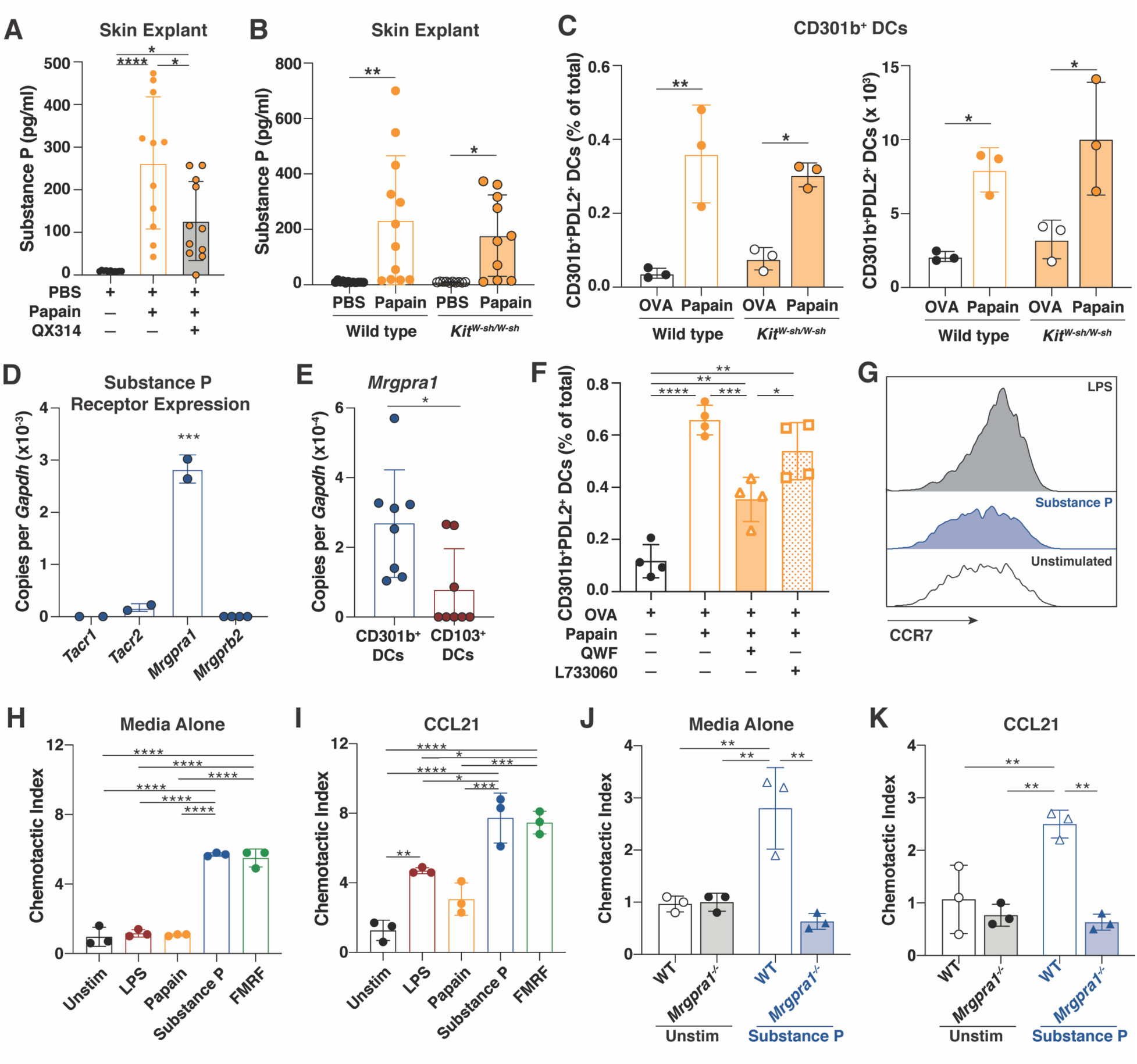
Substance P acts independently of mast cells to promote Th2-skewing CD301b^+^ dendritic cell migration through its receptor Mrgpra1. (**A**) Wild type mice were intradermally (i.d.) injected with PBS & papain with or without 1% QX314. Skin explants of the injected site were harvested and incubated in serum free media, and supernatant was tested by ELISA for Substance P. (**B**) Wild type or mast cell deficient *Kit^w-sh/w-sh^* mice were i.d. injected with PBS and/or papain. Substance P release from dermal explants was determined as in (A). (**C**) Wild type or mast cell deficient *Kit^w-sh/w-sh^* mice were i.d. injected with OVA or OVA/Papain and the dLN was harvested 24 hours later for flow cytometry analysis. The percent of CD11c^+^CD301b^+^PDL2^+^ cells (CD301b^+^PDL2^+^ DCs) out of total live cells in the dLN and the absolute number of activated CD301b^+^PDL2^+^ DCs were measured. (**D**) qPCR analysis of the Substance P receptors *Tacr1*, *Tacr2*, *Mrgpra1* and *Mrgprb2* on unstimulated BMDCs. (**E**) Wild type mice were immunized i.d. with papain and the dLN was harvested 24 hours later for flow sorting of live CD11c^+^CD301b^+^ (CD301b^+^ DCs) or CD11c^+^CD103^+^ (CD103^+^ DCs) cells. qPCR analysis of *Mrpgra1* expression on these sorted populations was performed. (**F**) Wild type mice were i.d. immunized with the indicated combinations of OVA, papain, QWF (Tacr1/Tacr2/Mrgpra1 inhibitor), and L733060 (Tacr1/Tacr2 inhibitor). The dLN was harvested 24 hours later for flow cytometry of CD301b^+^PDL2^+^ DCs shown as a percentage of total live cells in the dLN. (**G**) CCR7 expression by flow cytometry of CD11c^+^CD301b^+^ BMDCs left unstimulated (black unfilled histogram) or stimulated overnight with Substance P (blue filled histogram) or LPS (black filled histogram). (**H**) Wild type BMDCs were left unstimulated or stimulated overnight with LPS, papain, Substance P or FMRF (MRGPRA1 agonist), then washed, and replated on a Transwell membrane. Spontaneous migration in absence of an exogenous chemokine gradient (media alone) was evaluated and normalized to unstimulated BMDCs (chemotactic index). (**I**) Migration of BMDCs stimulated as in (H) across a Transwell membrane to CCL21 (100 ng/ml). (**J**) Migration in the absence of chemokines of unstimulated or Substance P stimulated wild type or *Mrgra1*^-/-^ BMDCs across a Transwell as in (H). (**K**) Migration to CCL21 of unstimulated or Substance P stimulated wild type or *Mrgra1*^-/-^ BMDCs across a Transwell as in (I). Symbols represent individual replicates (A, B, D, E, H-K) or mice (C, F). Histograms are representative individual samples (G). Bars indicate mean and error bars indicate SEM. Statistical tests: Ordinary one-way ANOVA with multiple comparisons (A, C-H, N), unpaired t test (B, K-M). * p<0.05, ** p<0.01, *** p<0.001, **** p<0.0001. Data are representative of at least two independent experiments, combined in (A, B, E) with each experiment including 2-5 mice per group.

### Substance P acts through MRGPRA1 on CD301b^+^ DCs to promote their migration

Substance P is a cationic neuropeptide that binds to its classical receptors TACR1 and TACR2 as well as the Mas-related G protein coupled receptors MRGPRA1 and MRGPRB2 in the mouse (Azimi et al., 2017; Azimi et al., 2016; McNeil et al., 2015). To investigate whether CD301b^+^ DCs expressed a Substance P receptor, we performed QPCR of bulk BMDCs, which are largely composed of the cDC2 population responsible for Th2 differentiation (Gao et al., 2013). BMDCs expressed *Mrgpra1*, but not other known receptors for Substance P (Fig. 6 D). To verify this expression on in vivo CD301b^+^ DCs, we sorted CD301b^+^ DCs and CD103^+^ DCs from the dLN after papain immunization. CD301b^+^ DCs, but not CD103^+^ DCs, expressed *Mrgpra1* indicating that CD301b^+^ DCs may be specifically sensitive to the effects of Substance P (Fig. 6 E). In order to determine the role of MRGRPA1, we sought to block its function using chemical antagonists. Specific antagonists of MRGPRA1 have not been described, so to determine the effects of MRGPRA1 blockade we compared the effect of the peptide antagonist QWF which blocks TACR1, TACR2, MRGPRB2, and MRGPRA1 to the small molecular antagonist L733060 which only blocks TACR1, TACR2, and MRGPRB2 (Azimi et al., 2017; Azimi et al., 2016). Consistent with a role for MRGPRA1 in CD301b^+^ DC migration, co-immunization with QWF, but not L733060, led to a defect in papain-induced CD301b^+^ DC migration (Fig. 6 F).

To this point, our data suggest that Substance P release from allergen-sensing TRPV1^+^ peptidergic nociceptive neurons activates local CD301b^+^ DCs through MRGPRA1 to initiate CD301b^+^ DC migration to the dLN. CD301b^+^ DCs require CCR7 to enter the draining lymphatics and then CCR7 and CCR8 to cross the subcapsular sinus and enter the dLN parenchyma (Sokol et al., 2018). Because CD301b^+^ DCs do not upregulate CCR7 expression upon allergen immunization, it has remained unclear how CD301b^+^ DCs are signaled to leave the skin parenchyma to migrate to the draining lymphatic vessels (Sokol et al., 2018). Unlike LPS, in vitro stimulation with Substance P had no effect on CCR7 expression (Fig. 6G). However, we found that stimulation with Substance P or the MRGPRA1 agonist FMRF induced DC chemokinesis across a Transwell membrane in the absence of any chemokine gradient (Fig. 6H) (Liu et al., 2009). Furthermore, despite their low level of CCR7 expression (Fig. 6G), BMDCs stimulated with Substance P or FMRF exhibited augmented migration to the CCR7 ligand CCL21 (Fig. 6I). The effect of Substance P on chemokinesis and chemotaxis was lost in BMDCs from *Mrgpra1^-/-^* mice, indicating that Substance P promotes DC migration through MRGPRA1 (Fig. 6J and K) (Meixiong et al., 2019b).

### TRPV1^+^ neurons are required for Th2 differentiation in response to papain

CD301b^+^ DCs are necessary for Th2-differentiation in response to cutaneous allergens and disruption of their migration into the parenchyma of the dLN blocks Th2-differentiation in response to papain (Kumamoto et al., 2013; Sokol et al., 2018). Consistent with their role in allergen-induced CD301b^+^ DC migration, we found that depletion of *Trpv1*^+^ neurons led to decreased allergen-induced Th2 differentiation (Fig. 7A and B). CD4^+^ T cells expressed significantly less IL-4 (Fig. 7C) and IL-13 (Fig. 7D), indicating a block in Th2 differentiation. There was no concurrent increase in background levels of IFNγ production by CD4^+^ T cells, suggesting that this block was at the level of CD4^+^ T cell activation and not simply Th2 polarization (Fig. 7E). Similarly, using IL-4-eGFP mice (4get) we found that co-immunization of OVA and papain together with QX314 or QWF inhibited the production of CD4^+^IL-4-eGFP^+^ cells as compared to OVA and papain alone (Fig. 7F) (Mohrs et al., 2001). Finally, although Substance P-induced DC migration was necessary for Th2-differentiation, it was not sufficient, indicating that an additional signal or cell is required to promote Th2-differentiation in vivo (Fig. 7F).

**Figure 7.**
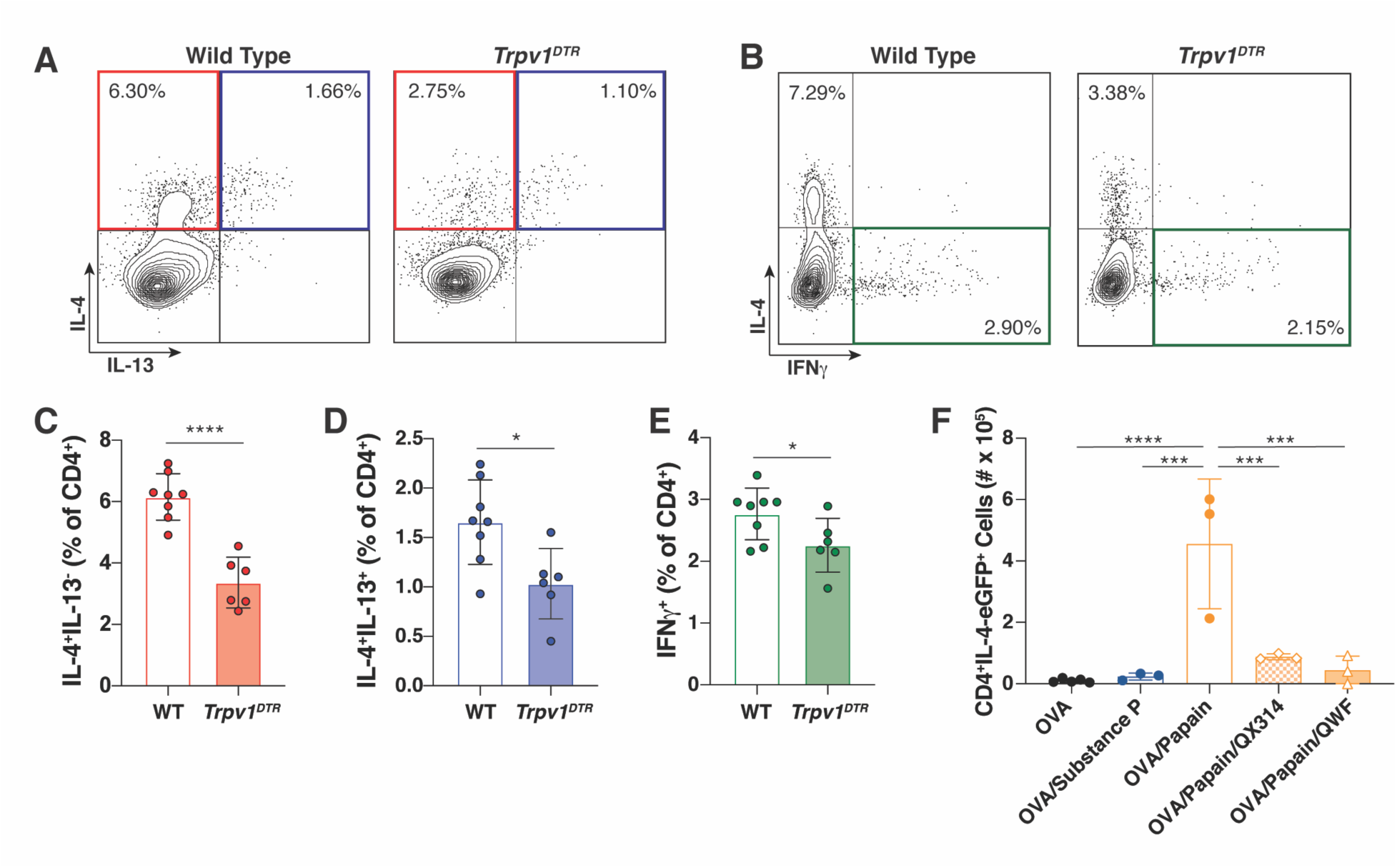
TRPV1^+^ neurons are required for Th2 differentiation in response to the protease allergen papain. (**A** and **B**) Wild type or *Trpv1^DTR^* mice treated with DT for 21 days and rested for 7 days were immunized with OVA/papain, with the dLNs being harvested 5 days later. Flow cytometry for IL-4 and IL-13 (A) and IL-4 and IFNγ (B) was performed after in vitro restimulation with PMA/Ionomycin. (**C-E**) Percent of CD4^+^ cells gated as in (A and B) that are IL-4^+^IL-13^-^ (C), IL4^+^IL-13^+^ (D), or IFNγ^+^ (E). (**F**) 4get (*IL- 4^eGFP^*) mice were immunized with OVA, OVA/Substance P, OVA/Papain, OVA/Papain/1% QX314, or OVA/Papain/QWF, with the dLN being examined 4 days later for total number of CD4^+^IL-4-eGFP^+^ cells. Symbols represent individual mice (C-F). Bars indicate mean and error bars indicate SEM. Statistical tests: Ordinary one-way ANOVA with multiple comparisons (F), unpaired t test (C-E). * p<0.05, ** p<0.01, *** p<0.001, **** p<0.0001. Data are representative of at least two independent experiments, with each experiment including 2-5 mice per group.

## DISCUSSION

In the classical PAMP-PRR signaling paradigm, tissue DCs act as direct sensors of Type-1 immunogens such as bacteria and viruses to initiate adaptive immune responses (Iwasaki and Medzhitov, 2015). Here we show that this paradigm may not be applicable to Type-2 (allergic) immunity. We found that TRPV1^+^ neurons act as the primary allergen sensor, eliciting the sensation of itch and pain and the release of Substance P upon exposure. Substance P then stimulates proximally located Th2-skewing CD301b^+^ DCs through MRGPRA1 to promote their movement. Activation of sensory neurons is necessary for the initiation of Th2 differentiation. We propose that DCs do not directly sense allergens and instead require the initial activation of a primary sensor. In the case of proteolytically active allergens such as papain, HDM, or Alternaria, the sensory nervous system acts as that primary sensor of allergen exposure to initiate the adaptive Type-2 immune response. IL-4 and IL-13, produced in the context of chronic allergic inflammation can decrease the activation threshold of itch sensing neurons, which could link allergen-induced acute activation of the sensory nervous system with chronic allergic inflammation (Oetjen et al., 2017).

Our study reveals that Substance P can directly induce the migration of CD301b^+^ DCs, of the cDC2 lineage, to the dLN. Importantly, Substance P induced migration is not sufficient for Th2-differentiation. This is consistent with studies showing an important role for innate alarmins in Th2-differentiation and allergic inflammation, but only a partial role in DC migration (Besnard et al., 2011; Cayrol et al., 2018; Palm et al., 2013). Both signals may be required to fully license the DC to promote Th2-differentiation, or they may individually act on DCs and putative accessory cells that provide the necessary skewing information for Th2-differentiation (Halim et al., 2018). This two-signal requirement could be essential to reduce the risk of non-specific or bystander activation of DCs. Unlike PAMPs, which DCs normally interact with in the context of antigens, indirect activation of DCs by neurons separates the activation trigger from the antigen. If both sensory neuron activation and alarmins are required for full DC licensing, this may limit DC activation to those DCs in areas of high antigen concentration. This would increase the likelihood that activated DCs are presenting the relevant antigen. However, this separation of antigen from adjuvant detection runs the risk of activating DCs presenting bystander antigens and could underlie the clinical observation of “allergen creep” in which atopic individuals progressively gain additional allergen sensitizations. Indeed, separation of antigen-adjuvant detection could explain why chronic low-level allergen exposure can lead to immune sensitization, why allergic individuals are often polysensitized, and also why atopic dermatitis patients with chronic itch are at higher risk of developing sensitization to food proteins found in household dust (Brough et al., 2015; Migueres et al., 2014).

Why is Substance P required for DC migration in the Type-2 immune response? Unlike in the PAMP-PRR signaling paradigm, DCs fail to directly sense allergen-associated molecular patterns and do not upregulate CCR7 upon allergen exposure. We propose that the Substance P signal is necessary to induce chemokinesis, allowing Th2-skewing CD301b^+^ DCs to release contacts from within the dermis. Unanchored CD301b^+^ DCs can then utilize their low-level expression of CCR7, which is necessary for their migration into draining lymphatics, to sense and respond to homeostatic CCL21 gradients and migrate to the dLN (Ohl et al., 2004; Sokol et al., 2018). This provides a potential mechanism by which allergen-activated DCs – in contrast to mast cells that can directly respond to allergens through IgE cross-linking – are able to sense and migrate out of the peripheral tissues without direct allergen detection and without concomitant upregulation of CCR7 (Lin et al., 2018).

Cysteine protease allergens, including papain and HDM, induce Ca^2+^ flux of DRG neurons in culture which indicates that nociceptive neurons directly sense allergens (Reddy et al., 2015; Serhan et al., 2019). How this sensing occurs is unclear. Protease activated receptors (PARs) 2 and 4 are cleaved by papain and have been associated with the itch response, but the interpretation of some of these studies is complicated by the fact that the PAR2 agonist SLIGRL also activates MRGPRC11 to mediate itch (Liu et al., 2011). Indeed, the MRGPR family plays a major role in neuronal itch sensing, with MRGPRA3 mediating chloroquine induced itch and MRGPRA1 mediating bilirubin induced itch (Liu et al., 2009; Meixiong et al., 2019b). MRGPRs are expressed on both mast cells and sensory neurons, permitting these cells to directly communicate through the release of preformed mediators including neuropeptides to promote itch and allergic inflammation (Meixiong et al., 2019a; Serhan et al., 2019). Here we find that CD301b^+^ DCs express MRGPRA1, but not other Substance P receptors, which allows these Th2-skewing DCs to sense Substance P from sensory neurons. This is in contrast to a report showing that BMDCs respond to Substance P by promoting Th1 differentiation indirectly after in vivo transfer by recruiting inflammatory DCs to the dLN (Janelsins et al., 2013). This response was ascribed to *Tacr1* expression on BMDCs based on their functional responsiveness to Substance P and a synthetic analogue. However, in the context of our study these data may be re-interpreted to be due to activation of MRGPRA1 on BMDCs. As shown in both of our studies, Substance P immunization alone is not sufficient to induce Th2 differentiation, though Substance P induced migration of “partially active” OVA-peptide loaded DCs to the dLN may lead to some background activation of T cells (Janelsins et al., 2013). This communication between the sensory nervous system and immune cells could be a rapid way to initiate protective host defenses (Cohen et al., 2019; Kashem et al., 2015), but its dysregulation could drive inflammatory or allergic diseases (Riol-Blanco et al., 2014; Talbot et al., 2015; Trankner et al., 2014).

The critical role of TRPV1^+^ sensory neurons and Substance P-driven DC migration in the development of allergic immune responses to cutaneous allergens provides a novel, targetable pathway for the prevention and treatment of allergic diseases. TRPV1^+^ neurons activated optogenetically or by *Candida* infection release CGRP that promotes IL-23 release from CD301b^+^ DCs and activation of the IL-17-mediated anti-fungal immune response (Cohen et al., 2019; Kashem et al., 2015). How *Candida* can promote CGRP release while protease allergens induce Substance P release is unclear. One possibility is that TRPV1^+^ neurons can differentiate between stimuli to promote different immune responses. Alternatively, given the heterogeneity in responsiveness to classic itch inducing ligands in papain-responsive neurons it is possible that *Candida* and protease allergens activate different subsets of TRPV1^+^ neurons. CGRP release may specifically induce innate IL-17 related inflammatory responses while Substance P may specifically induce initiation of the Type-2 adaptive immune response. The full extent of how CGRP and Substance P specifically activate different arms of Th-mediated immunity remain unclear, however neuropeptide specificity in initiating Type-1, Type-2, and Type-17 immune responses would allow for new and targeted therapeutics. Unlike topical corticosteroids that are currently the mainstay of cutaneous allergy therapeutics, targeting Substance P or its receptors would specifically blunt allergic responses while keeping other immune responses intact. Together, these data establish novel mechanisms for DC migration in allergic immune responses which may be applied to the prevention and treatment of allergic diseases.

## ACKNOWLEDGEMENTS

This work was supported by K08 AI121421 (C.L.S), DP2AT009499 (I.M.C.), R01AI130019 (I.M.C.), Center for the Study of Inflammatory Bowel Disease Pilot Feasibility Study Award DK043351 (C.L.S.), Massachusetts General Hospital Transformative Scholar Award (C.L.S.), AAAAI Foundation and Dr. Donald Y. M. Leung/JACI Editors Faculty Development Award (C.L.S.). Cytometric finding reported here were performed in the MGH Department of Pathology Flow and Image Cytometry Research Core, with support from the NIH Shared Instrumentation Program (1S10OD012027-01A1, 1S10OD016372-01, 1S10RR020936-01, and 1S10RR023440-01A1). We thank N. Andrew (MGH) for technical advice, M. Hoon (NIH) for providing *Trpv1^DTR^* mice, K. Blake and S. Sannajust (Harvard) for breeding *Tac1*^-/-^, *Nav1.8^cre^* and *tdTomato^loxSTOPlox^* mice, X. Dong (Johns Hopkins) for providing *Mrgpra1^-/-^* bone marrow.

## AUTHOR CONTRIBUTIONS

P.A.A., Z.N.A.D., and C.L.S. designed the study. P.A.A., Z.N.A.D., C.H.F., C.P., X.Z., T.V., R.B.C., O.A.C., and C.L.S. performed and/or analyzed experiments. P.A.A., Z.N.A.D., and C.L.S. wrote the manuscript. I.M.C. provided resources and advice. C.L.S. provided resources, reagents and funding. C.L.S. supervised the study.

## METHODS

### Contact for Reagent and Resource Sharing

Further information and requests for resources and reagents should be directed to and will be fulfilled by the Lead Contact, Caroline Sokol.

### Experimental Model and Subject Details

#### Mice

All animal experiments were approved by the Massachusetts General Hospital or Harvard Medical School Institutional Animal Care and Use Committee (IACUC). Mice were bred and maintained in a specific-pathogen-free (SPF) animal facility at Massachusetts General Hospital. C57BL/6 and *Kit^W-sh/W-sh^* mice were purchased from Charles River Laboratories (Wilmington, MA) or Jackson Laboratories (Bar Harbor, ME). Nav1.8^cre^ mice were originally from Dr. John Wood (University College London), and Trpv1^DTR^ from Dr. Mark Hoon (NIH). *Kaede* mice were crossed with *Trpv1^DTR^* mice to generate *Kaede x Trpv1^DTR^* mice. *Nav1.8^Cre^* mice were crossed with *tdTomato^loxSTOPlox^* to generate *Nav1.8^tdTomato^* mice. Bone marrow from *Mrgpra1^-/-^*mice in a C57BL/6 background was kindly provided by Dr. Xinzhong Dong (Johns Hopkins, MD). Mice were used in experiments at 5-14 weeks of age and age-matched littermates or purchased C57Bl/6 mice were used as wild type controls.

### Method Details

#### Immunizations

Mice were immunized intradermally (i.d) in the right side of the cheek (behavioral experiments), base of tail (Kaede experiments), or in footpads. As indicated, mice were immunized with 50 µg of papain, heat-inactivated papain, ovalbumin, histamine, bee venom phospholipase A_2_, or 40 µg of Capsaicin (all from Sigma-Aldrich); or 100 µg of Alternaria alternata (Greer). Where indicated, mice were injected with 10 µg LPS (InvivoGen), 5 µg CGRP (Sigma-Aldrich), 10 nmoles Substance P (Tocris) or 1% QX314 (Tocris). All dilutions were diluted in sterile 1x phosphate buffered saline, or PBS (Corning), except for capsaicin which was diluted in PBS and 6.4% DMSO. Where indicated, immunizations were performed with either 500 µmol of QWF or L733060 (both from Tocris) diluted in a 5% DMSO PBS solution.

#### Genetic Depletion of Trpv1+ Neurons

C57Bl/6, *Trpv1^DTR^*, Kaede, or Kaede x *Trpv1^DTR^* mice were injected intraperitoneally (i.p) with 0.2 µg of DT (Sigma-Aldrich) daily for 5 days then rested for 2 days repeated over a 21 day period. Mice rested for seven days post final DT injection. Effective depletion was confirmed through tail flick assay, and qPCR of DRGs. For the tail flick assay, the tail of each mouse was placed into 52°C water for a maximum of ten seconds, or less if mouse removed the tail from water.

#### Behavioral Analysis

For pain and itch assays, mice were allowed to acclimate to cage apparatus for two hours prior to filming, with food and water provided. A white noise machine (Marpac) was used to reduce distractions from behavioral response. After habituation, mice were immunized i.d. in the right cheek with allergen administered in a 25 µL vehicle under brief isoflurane anesthesia. Mice were videotaped in isolation for 20 minutes. Videos were then examined for quantification of wiping (single ipsilateral paw to injected cheek) and scratching (ipsilateral hind paw to injected cheek). Data was quantified in five-minute intervals.

#### Calcium Imaging of DRG Neuronal Cultures

DRGs were harvested and digested in a mix of collagenase (2.5 mg/ml) and dispase (1 mg/mL) for 80 min while shaking at 37°C, DRGs were then resuspended in DMEM/F12 (Thermo Fisher) and mechanically triturated using needles of different sizes (18G, 21G, 26G, six times each). The cell suspension was carefully plated on laminin precoated 35 mm plates in DMEM/F12 supplemented with NGF 25 ng/mL and GDNF 2 ng/mL and cultured overnight.

For calcium imaging, cells were loaded with 5 µM Fura-2-AM / DMEM/F12 (Thermo Fisher) at 37°C for 30 min, washed 3x, and imaged in 2 mL Krebs-Ringer solution (Boston Bioproducts) (KR: 120 mM NaCl, 5 mM KCl, 2 mM CaCl_2_, 1 mM MgCl_2_, 25 mM sodium bicarbonate, 5.5 mM HEPES, 1 mM D-glucose, pH 7.2 ± 0.15). 100 µg Papain or Vehicle (PBS) stimulation were followed by 100 µM histamine, 1 mM chloroquine, 1 µM capsaicin and 80 mM KCl sequentially application to identify shared neuronal responses of itch sensitizing stimulants with each other and TRPV1^+^ neurons. Images were acquired with alternating 340/380 nm excitation wavelengths, and fluorescence emission was captured using a Nikon Eclipse Ti inverted microscope and Zyla sCMOS camera (Andor). Ratiometric analysis of 340/380 signal intensities were processed, background corrected, and analyzed with NIS-elements software (Nikon) by drawing regions of interest (ROI) around individual cells as previously described (Lai, N.Y. et al. 2019). An increase from baseline greater than 15% was considered a positive response. The percentages of papain, histamine, chloroquine and capsaicin and vehicle responsive cells were quantified and plotted as proportion of KCI responsive cells. Response traces of single DRG neurons were generated using Microsoft Excel. Venn Diagram showing overlaps in response for the different stimulants was created using R studio 3.6.2. Violin plots were created using GraphPad PRISM software 8.4.

#### Microscopy and Image Quantification

Ears were harvested, split into dorsal and ventral halves, removed of hair (Veet) and placed on Flexigrid™ transparent adhesive film (Smith and Nephew Opsite) dermal side up. Fat was removed and each tissue sample was floated on 3 mL of DMEM (Corning) supplemented with 3 mg/mL Dispase II (Sigma-Aldrich) per well of a 6-well plate, dermal side down, for 90 minutes at 37°C in 5% CO_2._ Afterwards, the samples were dried and the dermal tissue was gently peeled away from the epidermal tissue. Dermal slices were then isolated, rinsed in PBS, and fixed in a 4% paraformaldehyde (Electron Microscopy Sciences) PBS solution for 1 hour at 4 °C. After washing in PBS, samples were incubated overnight with a primary antibody solution of biotinylated anti-Tuj1 (BD Biosciences) diluted in PBS with 0.2% TritonX-100 (Sigma-Aldrich) and 10% heat inactivated goat serum (Jackson ImmunoResearch). Samples were washed and then stained with Streptavidin-AF488, CD301b-AF647, and MHCII-BV421 (all from BioLegend) diluted in PBS/0.2% TritonX-100 for 1 hour at room temperature with shaking. Washed ears were mounted using Prolong Diamond Antifade Mountant (Thermo Fisher Scientific) prior to imaging on a Zeiss LSM confocal microscope. Images were analyzed using Zen Blue to quantify the distance of MHCII^+^CD301b^+^ cells and MHCII^+^CD301b^-^ cells from their nearest Nav1.8^+^ neuron as measured in 3D Ortho.

#### Kaede Photoconversion

A 2 x 2 cm patch of skin above the base of tail was shaved followed by application of a chemical depilatory agent (Veet) for 1 min, then removed by multiple washings with PBS. The exposed skin was subjected to violet light (420 nm) using a Bluewave LED visible light curing unit (Dymax) with a 420 nm bandpass filter (Andover Corp). Skin was exposed to this wavelength with the light source at a maximum power, approximately 7.5 cm away from the skin for 5 min. Following photoconversion, mice were immunized subcutaneously in the shaved region as described above. 24 hr following immunization, dLNs were isolated and analyzed for skin DC migration using flow cytometry.

#### Flank Explant Assay

Wild type mice were shaved at the flanks and injected as described. Mice were immediately sacrificed and punch biopsies of the shaved flanks were collected and rapidly transferred into a 24-well plate containing 1mL of serum free DMEM. Explants were incubated for 30 min at 32 °C with gentle rotation (150 rpm) before supernatants were collected for analysis.

#### Enzyme-Linked Immunosorbent Assay (ELISA)

Mouse Substance P and CGRP were detected using competitive mouse Substance P and CGRP ELISA kits (both from Cayman Chemical). All procedures were carried out according to the manufacturer’s protocols. Samples were assayed on Softmax Pro Software in triplicates and concentrations were determined from a standard curve.

#### Flow Cytometry and Cell Sorting

Harvested lymph nodes were subjected to digestion at 37°C in DNase I (100 mg/mL, Roche), Collagenase P (200 mg/mL, Sigma-Aldrich), Dispase II (800 mg/mL, Sigma-Aldrich), and 1% fetal calf serum (FCS) in RPMI. At 8 min intervals, supernatant was removed and replaced with fresh enzyme media until no tissue fragments remained. Supernatant was added to added to stop buffer (RPMI/2mM EDTA (Gibco)/1% FBS) and filtered prior to antibody staining. Enzymatically digested cell suspensions were incubated for 15 min at 4°C in PBS with 0.5% FCS with the following antibodies: anti-CD16/CD32, anti-CD11b, anti-CD11c, anti-CD103, anti-CD301b, anti-CD4, and anti-PDL2 (all from BioLegend). Intracellular cytokine staining was performed after PMA/Ionomycin (Sigma) stimulation of single cell suspensions in the presence of GolgiPlug for 4 hours prior to intracellular staining using BD Cytofix/Cytoperm kit (BD Biosciences). Viability was determined using Fixable Viability Dye eFluor 780 (Invitrogen). Samples were run on Beckman Coulter’s CytoFLEX S, BD Fortessa X20 or BD FACSAria Fusion and analyzed using FlowJo (Version 10) (TreeStar).

#### Dorsal Root Ganglia Cultures and Stimulation

Dorsal root ganglia (DRGs) were harvested from mice (7-14 weeks old) and dissected into DMEM supplemented with 0.1% glucose, 0.1% L-glutamine, 0.1% sodium pyruvate (Corning) with added 10% HI FBS (Sigma-Aldrich) and 1% Pen/Strep (Lonza). DRGs were then dissociated in 1.25 mg/mL Collagenase A (Roche) + 2.5 mg/ml Dispase II (Sigma-Aldrich) in 1x PBS (Corning). Dissociated DRGs were washed in supplemented DMEM and were triturated with glass pipettes of decreasing sizes into 2 mL of fully supplemented DMEM and 62.5 U/mL Deoxyribonuclease I type IV (Sigma-Aldrich). The DRG cells were then passed over a 3% bovine serum albumin (BSA) (Sigma-Aldrich) gradient and resuspended in Neurobasal-A medium (Invitrogen) supplemented with B27 (Invitrogen), 1% GlutaMAX™, and 1% Pen/Strep, with added growth factors 0.01 mM arabinocytidine (Ara-C), 0.002 ng/µL glial derived neurotrophic factor (GDNF) (Sigma-Aldrich) and nerve growth factor 2.5S (NGF 2.5S) (Invitrogen). For culture in a 96-well sterile tissue-culture (TC) treated plate (Corning), around 12,000 DRG neurons/well were plated at a standard 30 µL/well into 10 μg/ml laminin (Sigma-Aldrich) pre-coated plates and incubated for one hour at 37°C in a CO_2_ tissue incubator (Thermo Scientific). After the one-hour incubation, 100 µL of supplemented Neurobasal-A medium with added growth factors was added to each well. The DRGs incubated for 48 hours, with daily feeding with fresh neurobasal-A medium with added growth factors. For stimulations, media was replaced with 200 µL/well of supplemented Neurobasal-A medium without the addition of growth factors and incubated for 30 minutes. The DRG cultures were either left unstimulated with PBS or stimulated with papain, heat-inactivated papain, increasing concentrations (100 µg/mL, 150 µg/mL, or 200 µg/mL) of Alternaria extract or house dust mite extract.

#### Quantitative PCR (qPCR)

Total RNA from sorted cells, DRGs, or BMDCs was isolated using the QIAGEN RNeasy Micro Kit in accordance with the manufacturer’s protocol. 50 µL of cDNA per reaction was generated using random hexamers, Oligo (dT), magnesium chloride, dNTPs (10 mM), Reverse Transcriptase, and RNase inhibitor (all from Thermo Fisher Scientific). A Roche LightCycler 96 Real-Time PCR System was utilized to quantify gene expression, with SYBR Green Master Mix (Roche). The reaction cycles were: 95°C for 600 s, then 95°C, 60°C, and 72°C for 10 s each for 45 total cycles, followed by 95°C for 10 s, 65°C for 60 s, and finally 97°C for 1 s. Fluorescence was quantified during each amplification. Quantification cycle (Cq) values for each sample were determined using Roche LightCycler 96 Software 1.1. Microsoft Excel was used to determine copy values for Cq in order to divide by *Gapdh* copy values for ratio analysis.

#### Generation of Bone Marrow-derived Dendritic Cells (BMDCs)

BM was harvested from 5- to 8-week-old mice by mechanical disruption using a mortar and pestle, followed by incubation with red blood cell lysis buffer (Sigma-Aldrich). BM cells were cultured in growth media supplemented with 10% HI FBS (Sigma-Aldrich), 1% Pen/Strep (Lonza), 1% GlutaMAX™(Gibco), 1% HEPES buffer (Corning), 1% Non-Essential Amino Acid Solution (Lonza), 1% Sodium Pyruvate (Gibco), 0.1% 2- mercaptoethanol (Gibco), with 20 ng/mL of granulocyte-macrophage colony-stimulating factor (GM-CSF) (Peprotech) at a density of 0.7 x 10^6^ cells/mL, fed on days 2 and 4 and stimulated on day 5 of culture.

#### Chemotaxis Assays

On day 5 of culture, BMDCs were stimulated with: LPS (100 ng/mL, InvivoGen), papain (100 µg/mL, Sigma-Aldrich), Substance P (0.5 µM, Sigma-Aldrich), Nle-Arg-Phe amide, or FMRF (30 µM, Sigma-Aldrich). Cells were then harvested. Chemotaxis was performed on these harvested cells using 5 µm pore size Transwell membranes (NeuroProbe). 32 µL of CCL21 (100 ng/mL) or media alone with 0.5% BSA in RPMI, were added to the wells of the lower chamber. The Transwell membrane was then outfitted to the microplate, after which 100,000 cells in 50 µL of chemotaxis media were added in triplicate. Plates were incubated for 2 hr at 37°C in 5% CO_2_. Afterwards, the membrane was quickly rinsed with sterile PBS and spun down for one minute at 1500 rpm to collect any migrated adherent cells. The membrane was then removed, and migrated cells were counted.

### Key Resources Table

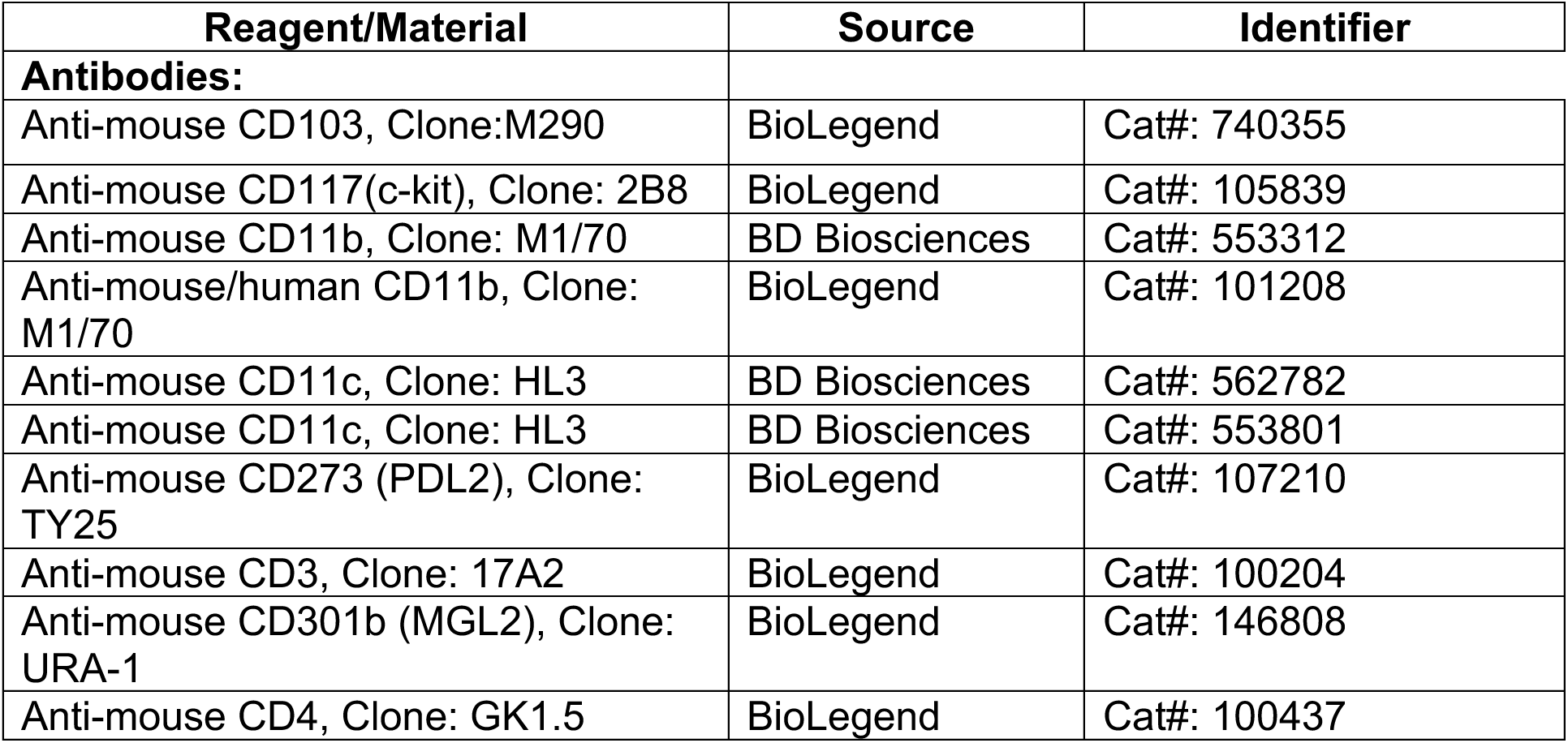
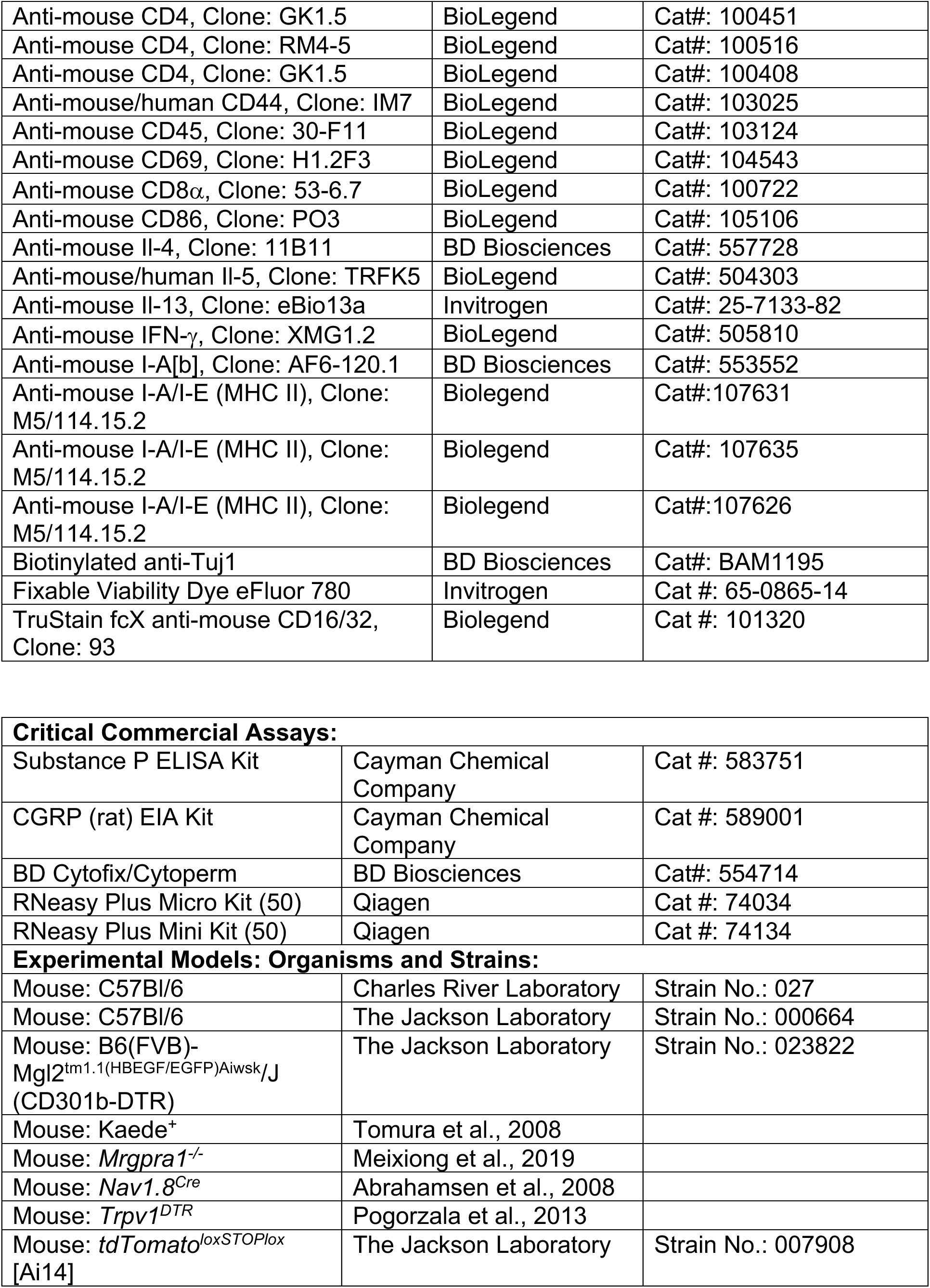
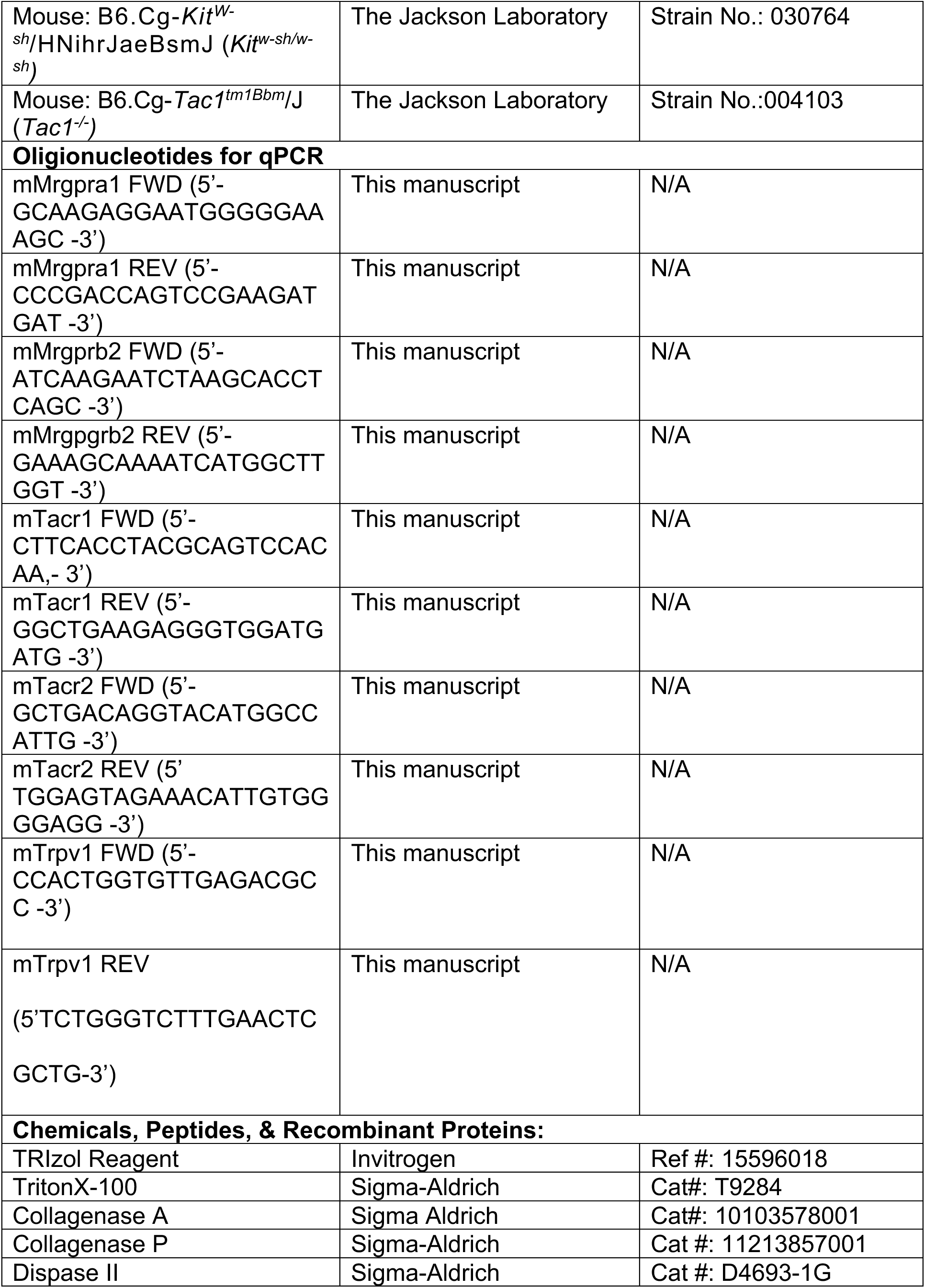
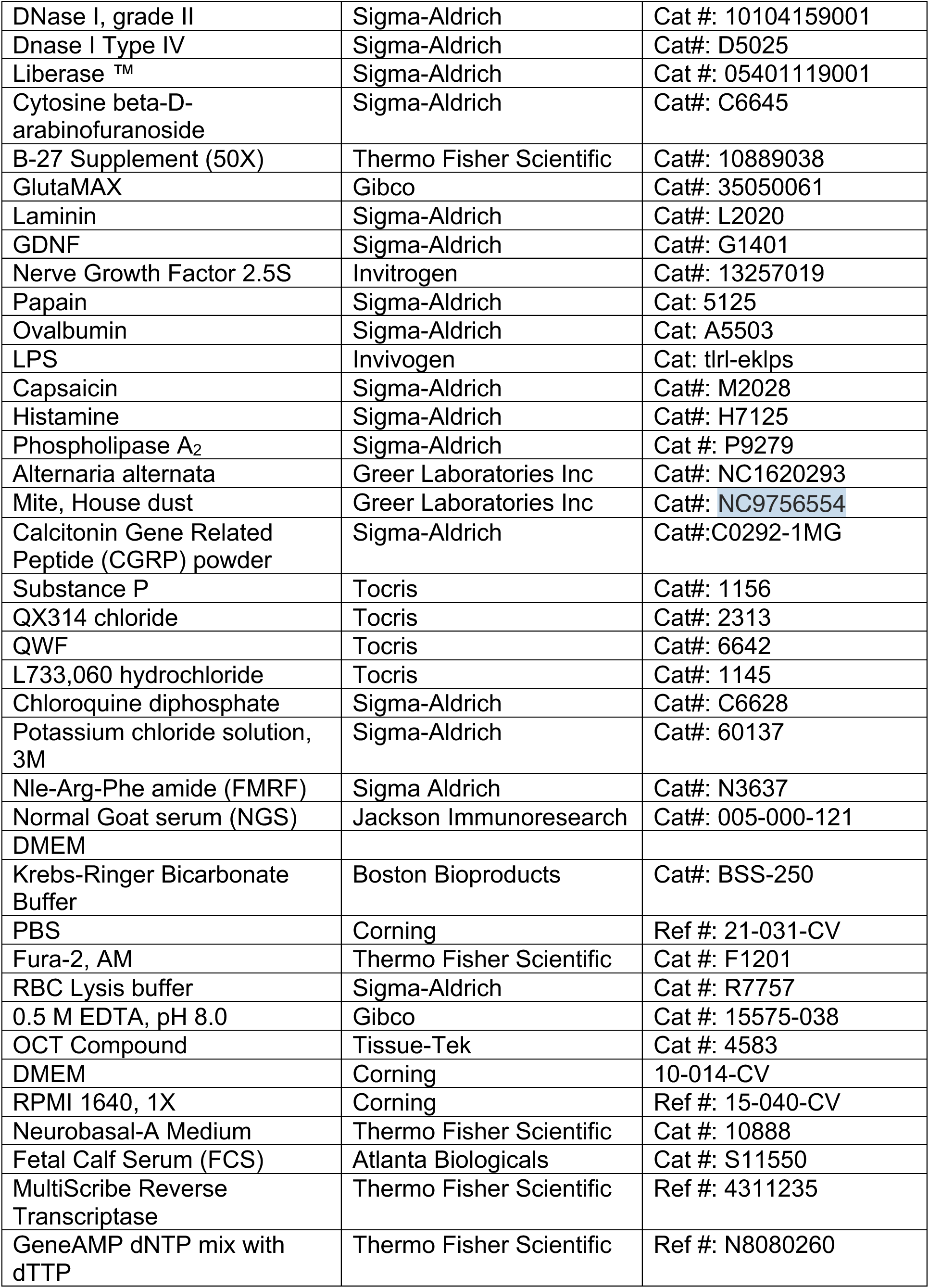
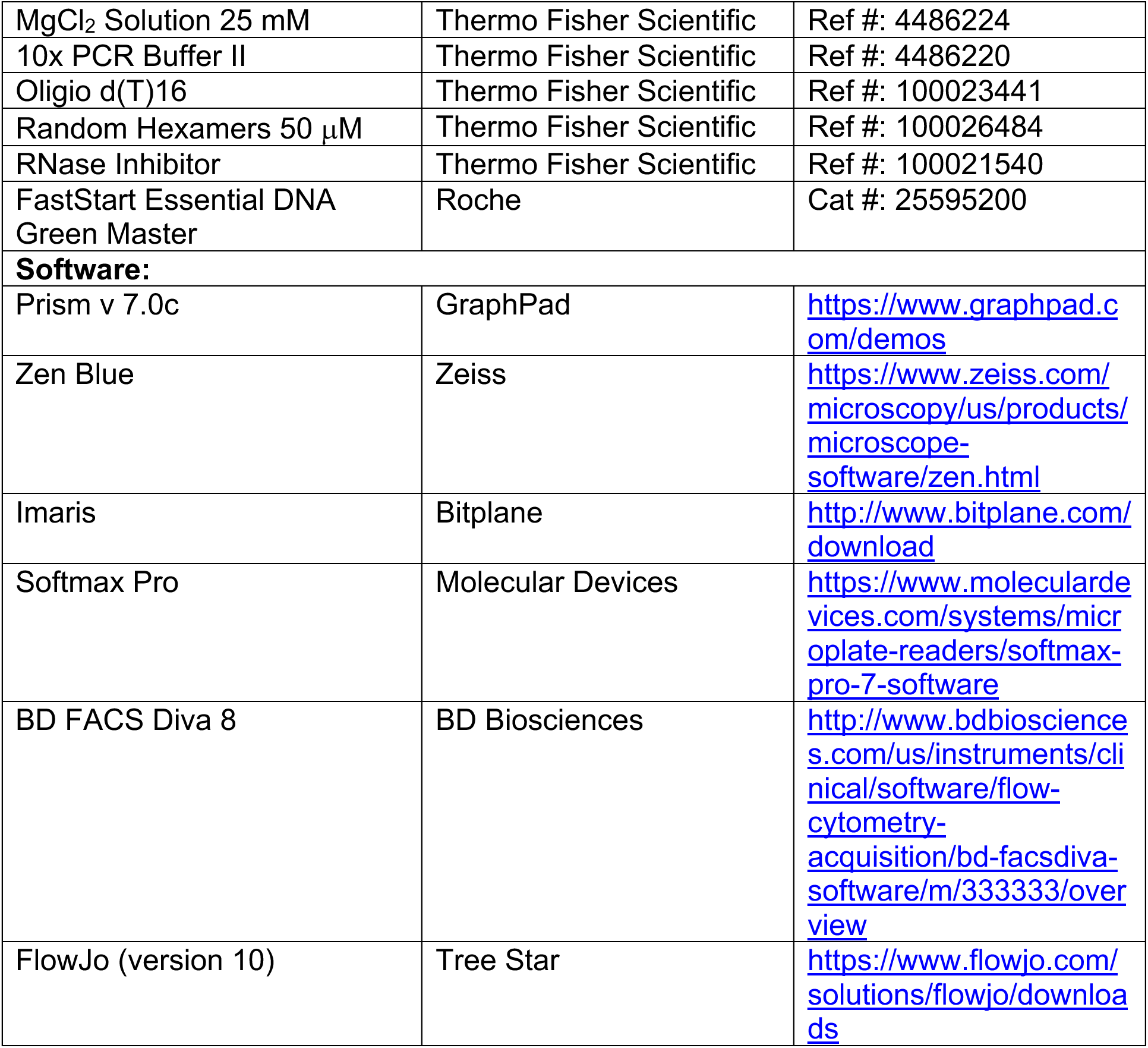

