## Supplementary Figure for "Allergen-induced dendritic cell migration is controlled through Substance P release by sensory neurons"

**SUPPLEMENTARY INFORMATION – Aderhold et al**

Figure S1: Aderhold et al

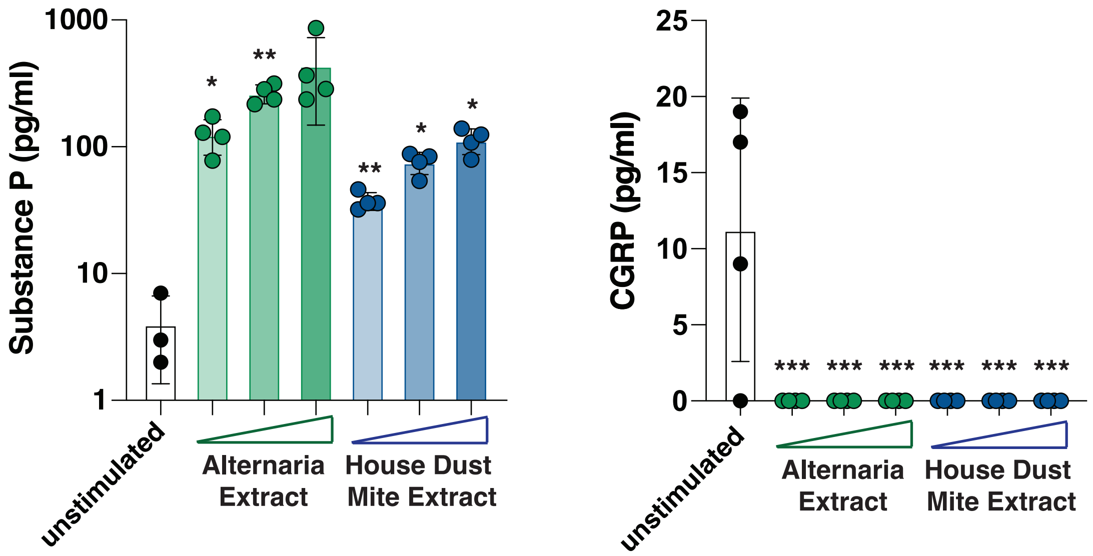


**Figure S1. Substance P release by other allergens.**

Dorsal root ganglia were harvested from wild type mice and were left unstimulated with PBS, or stimulated with increasing concentrations (100 µg/ml, 150 µg/ml or 200 µg/ml) of Alternaria extract or house dust mite extract. Supernatants were measured for Substance P and CGRP release by ELISA. Each dot represents one replicate. Statistical tests: Ordinary one-way ANOVA with multiple comparisons. Each bar compared to unstimulated control. * $p<0.05$, ** $p<0.01$, *** $p<0.001$. Data are representative of two independent experiments.
